# Lifespan trajectories, metabolic associations, and links to fluid intelligence of brain temporal dynamics characterized by Multiscale Entropy

**DOI:** 10.1101/2025.05.08.652915

**Authors:** Gianpaolo Del Mauro, Xinglin Zeng, Ze Wang

## Abstract

Brain entropy (BEN) offers a unique perspective into brain’s functioning by characterizing the irregularity and randomness of brain activity. We used multiscale entropy (MSE) to estimate BEN from rs-fMRI at different timescales (from Scale 1 to Scale 5). Using a GMALSS framework, we showed that the aging trajectory of entropy differs at different timescales. Specifically, entropy increases with aging at Scale 1, the curve flattens at Scale 2, and the pattern finally reverses from Scale 3 onward, showing that entropy decreases with aging. Entropy at coarser scales (from Scale 2) showed positive correlations with measures of brain metabolism, including oxygen and glucose metabolism, whereas Scale 1 entropy displayed a trend for negative correlation. In addition, we explored the relationship between BEN and fluid intelligence (FI). Using moderated mediation models, we showed that brain entropy mediates the association between age and FI, and that the relationship between entropy and FI is itself moderated by age, indicating that entropy can either attenuate or exacerbate age-related differences in cognitive performance, depending on life stage. Finally, we used BEN to predict age. The best accuracy was achieved using the GPR (R^2^=0.80, MAE=7.25 years). Importantly, the difference between predicted and chronological age (brain age gap, BAG) was weakly associated with FI in an age-dependent manner, suggesting that lower entropy patterns relative to age-matched individuals may be associated with better cognitive performance during midlife, whereas higher entropy patterns may be associated with better cognition in older age.

## Introduction

The notion of brain entropy (BEN) is used in neuroscience to characterize the irregularity or randomness of brain activity. In this sense, BEN is linked to Shannon’s Information Theory, which relates entropy of a time series to its information capacity (1). Several approximate metrics have been developed to estimate brain entropy using brain signals at a single voxel or regions of interest (2). Among these, Sample Entropy (SampEn) has been devised as a measure of complexity and regularity of clinical and experimental time series that can be applied to typically short and noisy data (3). In neuroimaging studies, SampEn can be applied to resting-state fMRI (rs-fMRI) time series to estimate the spontaneous regularity of BOLD signal (4). While we focus here on rs-fMRI data, entropy quantification has also found fertile application in techniques, such as EEG and MEG, that provide higher temporal resolution.

Previous studies have explored the age trajectories of BEN, with different studies reporting that BEN increases (5–7) or declines (8–11) with age. One possible explanation for these discrepant findings is that BEN–age associations depend on the timescale considered, with finer timescales showing increases and coarser timescales showing decreases in normal aging (5). This pattern suggests that entropy measured at different timescales may capture distinct neurophysiological processes. Specifically, finer timescales may be more sensitive to fast, local fluctuations in neural activity, whereas coarser timescales may reflect slower, more distributed dynamics. Accordingly, age-related changes in BEN may not be uniform, but instead reflect differential alterations in these dynamics across timescales.

Entropy at different timescales can be easily quantified using an extension of SampEn, namely the Multiscale Entropy (MSE) (6, 7). Using SampEn, entropy increases with the degree of signal irregularity and is maximum for completely random systems. Significantly, electrophysiological data suggest that aging is accompanied by an increase in spontaneous, noisy baseline neural activity (8). Hence, higher SampEn in aging can potentially reflect increased randomness of the brain signal rather than increased complexity. By characterizing entropy at different time scales, MSE has been used as an approximate approach for assessing signal complexity (6, 7). Recently, the normative age trajectory (9) of MSE was assessed and showed an inverted U-shaped trajectory, where the complexity of the brain signal initially increased, peaking at approximately age 40, and then declined (10). However, this study used a coarse-grained brain parcellation to estimate entropy and calculated the average entropy across timescales, thereby potentially overlooking differences in the trajectories of different timescales. In this study, we aim to overcome these limitations by estimating normative aging and developmental trajectories of BEN in a large lifespan cohort using a voxel-wise approach, while modeling entropy independently at each timescale. To further characterize the neurophysiological relevance of MSE, we examined associations of entropy at different timescales with markers of brain metabolism, including cerebral blood flow (CBF), oxygen metabolism, and glucose metabolism. We predict that different timescales will display distinct age trajectories, and specifically that entropy will increase across the lifespan at finer scales and decrease at coarser scales. In addition, we hypothesize that entropy at different timescales will display differential associations with brain metabolism.

Consider citing more BEN-related papers from us and Kay Jann or Danny Wang’s group: rsfMRI-derived BEN (4, 11–15) has been shown to be reproducible, mostly independent of several other more established functional brain measures (16), self-organized (4, 12–17), correlated with biological variables (17–19), education years (18), cognitive functions and task activations (17, 20, 21), sensitive to several disease conditions (22–31), and importantly sensitive to stimulant or medicine or hormone or neuromodulation (32–36). In particular, BEN has been associated to general cognitive ability and specifically to fluid intelligence (FI), with FI increasing at lower BEN (18). However, if the aging trajectories of entropy diverge from finer to coarser timescales, as hypothesized, then the functional meaning of entropy, and hence its relationship to intelligence, may also change during the lifespan. Using moderated mediation models (39), we propose that the relationship between age and FI will be mediated by changes in BEN, and that age will in turn moderate the relationship between BEN and FI. Moreover, we predict that these relationships will be different across timescales. Further, by means of machine learning, we will use MSE to predict age and compute the brain age gap (GAP), defined as the difference between the predicted and the chronological age. We hypothesize that the BAG will be correlated to FI.

## Materials and methods

### Datasets and Participants

Three datasets from the Human Connectome Project (HCP) were aggregated: HCP-Development (HCP-D) (40), HCP-Young Adults (HCP-YA) (41), and HCP-Aging/Aging Adult Brain Connectome (AABC) (40). As the precise age of participants with 90 years or more was not provided, these participants were excluded from the sample. The final sample used in this study included 2776 participants from 8 to 89 years old.

### rs-fMRI data

HCP provided fully preprocessed rs-fMRI data (40, 42), each participant including up to four rs-fMRI runs acquired over two days. Runs acquired on the same day used opposite encoding directions (HCP-YA: L-R/R-L; HCP-D and AABC: A-P/P-A). Voxel-wise time series were extracted using a brain mask, followed by spatial smoothing (Gaussian kernel, FWHM = 2.5 mm). The data were then linearly detrended and temporally high-pass filtered at 0.008 Hz. Finally, runs acquired on the same day were concatenated to maximize the time series length (HCP-YA: 2,400 time-points; HCP-D and AABC: 956 time-points). Two final rs-fMRI time series, named Rest1 and Rest2, were obtained. Framewise displacement (FD) was calculated separately for each run and then averaged across runs acquired on the same day, corresponding to the concatenated time series. Participants with an average FD > 0.3 were excluded from the sample.

### MSE mapping

SampEn is calculated as the logarithmic likelihood that a small section (within a window of a length *m*) of the data that “matches” other sections will still “match” the others if the section window length increases by 1 (3). A “match” is identified when the distance between two compared time segments is smaller than a threshold *r*.

Denote a rs-fMRI time-series of a voxel as *x* = [*x*_1_, *x*_2_, … *x_N_*], where *N* is the number of time points. SampEn starts with forming a series of vectors, called embedded vectors, each with *m* consecutive points extracted from x: *u_i_* = [*x_i_*, *x_i_*_+1_, … *x_i_*_+*m*−1_], where *i* = 1, …, *N* − *m* + 1. Using the pre-specified distance threshold 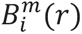 counts the number of *u_i_* whose Chebyshev distances to *u*_j_ (*i* = 1, *to N* − *m*, *and j* ≠ *i*) are less than *r*, so does 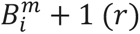 for the dimension of *m* + 1. By averaging across all possible vectors, it is obtained:

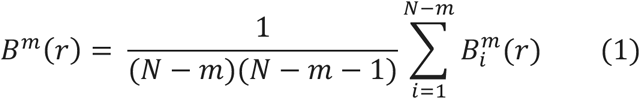

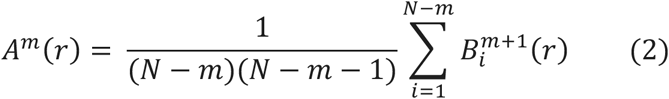

And the SampEn is calculated as:

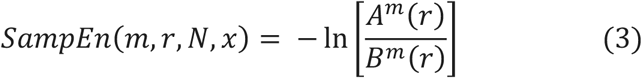

MSE is a combination of coarse-graining and SampEn: after down scaling the original time series to a specific scale, the single-scale SampEn is calculated (6, 7). Coarse graining is a moving average process using non-overlapping windows. For a given time series *x* = [*x*_1_, *x*_2_, … *x_N_*] and a scale s, averaged timepoint 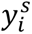 of the new time series is given by

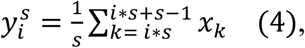

and the length of the new time series is 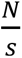

Voxel-wise MSE was computed from Rest1 and Rest2 using s =1-5, *m* = 2, and *r* = 0.3 (10). At each scale, average entropy was calculated using a gray matter mask derived from the cortical Schaefer 200 atlas and the subcortical Tian S2 atlas (43, 44). To assess the potential influence of head motion on BEN estimates, we computed Pearson’s correlations between average FD and BEN at each scale. Results indicated low-to-moderate correlations, ranging from *r*=0.07 at Scale 1 to approximately *r*=−0.24 at Scale 4. Negative correlations remained stable from Scale 2 to Scale 5.

Regional-level BEN was additionally obtained by averaging entropy values within the areas of above-mentioned brain atlases, including in total 232 regions-of-interest (ROI) A workflow chart of the analyses is presented in Figure 1

**Figure 1.**
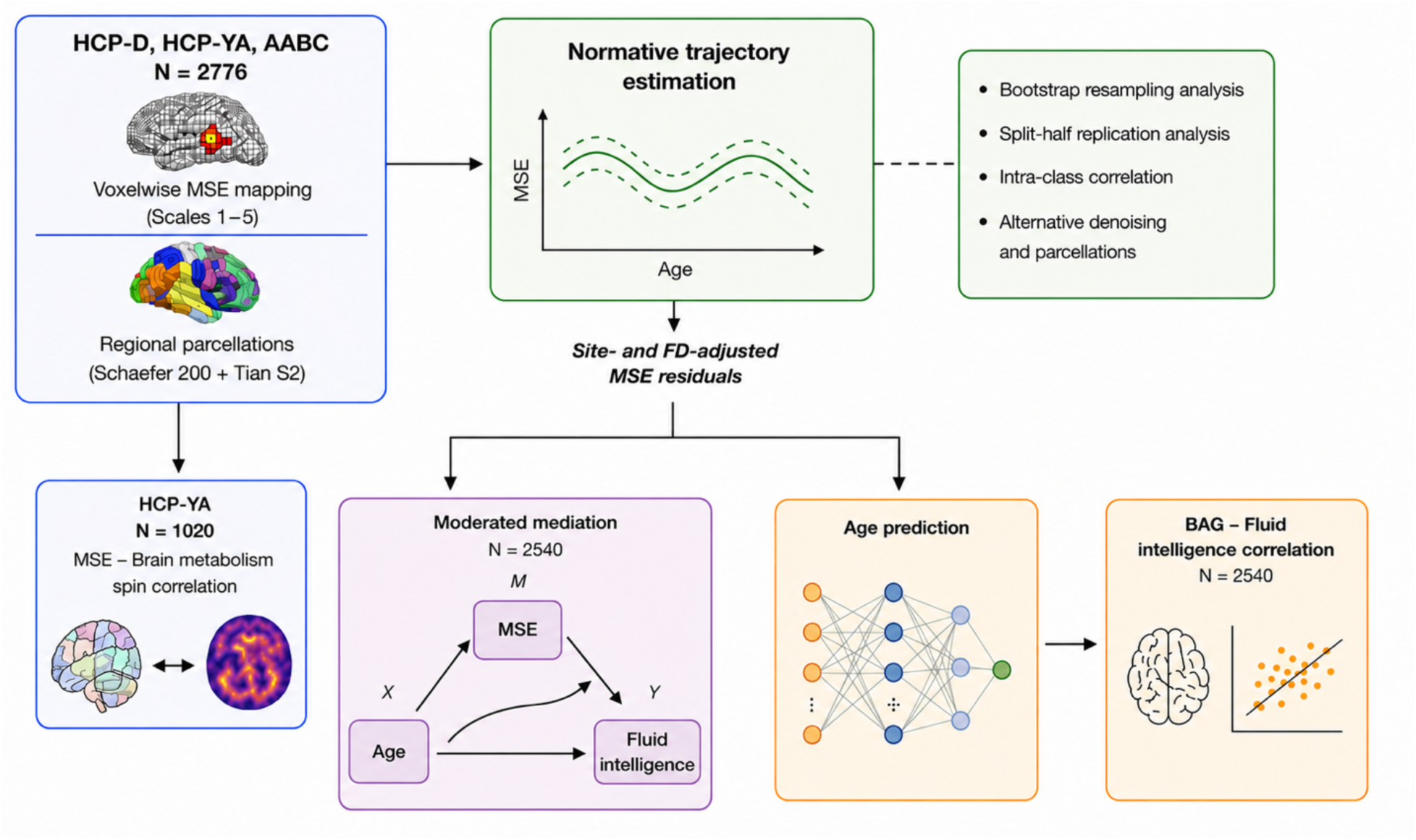
Workflow chart of the analyses. Multiscale Entropy (MSE) from Scale 1 to Scale 5 was estimated voxelwise using the Human Connectome project (HCP) development (D), young-adult (YA), and Aging Adult Brain Connectome (AABC) data. Entropy values were averaged inside cortical and subcortical areas using the Schaefer 200 and Tian S2 parcellations. HCP-YA data were separately used to test the association between brain entropy and brain metabolic metrics using spin correlation. Normative trajectory was estimated independently for each scale using the GAMLSS framework. Secondary and sensitivity analysis included: bootstrap resampling analysis, split-half replication analysis, intra-class correlation, and alternative denoising and parcellations. Site- and framewise displacement (FD)-adjusted entropy values were extracted from the GAMLSS models and used for (i) moderated mediation models, where Age was the independent variable (X) and moderator, entropy the mediator (M), and fluid intelligence the dependent variable (Y); and (ii) age prediction using machine learning. The best-performing model was used to predict age and calculate the brain age gap (BAG), which was finally correlated with fluid intelligence.

### Modeling normative growth curves across the lifespan

#### (I) GAMLSS framework

To estimate normative growth patterns for BEN across cohorts at each scale, Generalized Additive Models for Location, Scale, and Shape (GAMLSS) were applied using the GAMLSS package (version 5.4.22) in R 3.6.3 (45). Model fitting involved selection of an appropriate distribution family and specification of parameter models for the metric of interest. Metric-specific GAMLSS models were used to generate nonlinear normative growth curves and their first derivatives.

The fit of 29 continuous distribution families was evaluated separately for the average BEN of each timescale. Model performance was assessed using the Bayesian Information Criterion (BIC) (46), with lower values indicating better fit. Each distribution is parameterized by up to four parameters: location (μ), scale (σ), skewness (ν), and kurtosis (τ). Among the candidate distributions, the Gumbel (GU) distribution (μ, σ) provided the best fit for Scale 1; Johnson’s SU (JSU) (μ, σ, ν, τ) for Scale 2; Skew t Type 4 (ST4) (μ, σ, ν, τ) for Scale 3; t-family (TF) (μ, σ, ν) for Scale 4; and Skew t Type 3 (ST3) (μ, σ, ν, τ) for Scale 5.

In the final models, Age, Sex, and Site were included as predictors for μ and σ. Average FD was additionally added as predictor for μ. Only intercept was specified for ν and τ when these parameters were estimated by the distribution. Age was modeled using either a linear term or a spline function in the μ equation (47), with the degrees of freedom selected based on best model fit for each scale. For Scale 2, a linear age term provided the best fit. Sex and Site (HCP-Y and HCP-DA, the latter including HCP-D and AABC) were modeled as fixed effects. The rationale for treating HCP-D and AABC as a single site is that these datasets were acquired using the same acquisition parameters (40), and a previous study estimating normative CBF trajectories using HCP-D and HCP-A did not include site as a covariate in the GAMLSS model (48). Moreover, preliminary analyses in our data suggested that, regardless of whether HCP-D and AABC were modeled as separate sites or as a single site, site correction primarily affected the HCP-YA dataset. At the same time, combining HCP-D and AABC reduced multicollinearity between Age and Site. The final models estimated for each timescale are reported in the Supplementary Information (SI).

Normative developmental and aging trajectories were characterized by estimating age-dependent changes in the distribution of BEN, from which centile curves (e.g., 2.5th, 50th, and 97.5th percentiles) were derived. Centiles were obtained from the fitted GAMLSS model by computing the corresponding quantiles of the estimated distribution at each age.

The same GAMLSS models were computed for 232 ROIs to show the developmental and aging trajectories at the regional level.

From both whole brain and regional level models, site- and FD-adjusted residual entropy values were saved and used for subsequent analyses except correlations with brain metabolism.

#### (II) Sensitivity analyses

To validate the lifespan normative growth patterns, two sensitivity analyses were conducted: bootstrap resampling analysis and split-half replication analysis (49).

##### Bootstrap resampling analysis

To assess the robustness of lifespan growth curves, we conducted 1,000 bootstrap repetitions with replacement sampling. The sampling preserved the age, sex, and site proportionality of the original cohort by stratifying the lifespan based on ten age intervals (from 8 to 89 years). For each entropy scale, a growth curve was estimated in each bootstrap sample, yielding 1,000 bootstrap estimates of the 50th centile curve. The final median curve was obtained as the mean bootstrap estimate, and 95% bootstrap confidence intervals (CIs) were calculated as the 2.5th and 97.5th percentiles of the bootstrap distribution at each age.

##### Split-half replication analysis

To evaluate model reproducibility and generalizability, we performed a repeated bidirectional split-half validation. In each iteration, the dataset was randomly divided into two equal halves stratified by site. One half was used to train the GAMLSS model, and the fitted model was then applied to the held-out half to assess out-of-sample predictive performance. The procedure was subsequently repeated with the training and testing sets reversed.

Model generalizability was quantified using R-squared (R²), calculated from predictions in the held-out test set. Distributional calibration was assessed using quantile randomized residuals (randomized z-scores) derived from the fitted training models. Residual normality was evaluated using the Shapiro–Wilk statistic, with values approaching 1 indicating good agreement with the expected normal distribution. Additional calibration metrics included residual skewness and excess kurtosis, with values close to 0 indicating symmetric residual distributions and minimal deviations from normal tail behavior, respectively.

The entire split-half validation procedure was repeated 1,000 times. For each metric, the median and empirical 95% confidence interval (2.5th–97.5th percentiles) were calculated across the 1,000 repetitions.

### Intraclass-correlation

While the primary normative trajectory was estimated using average BEN from Rest1, centile scores for Rest2 were obtained by applying the GAMLSS model fitted on Rest1 to the corresponding Rest2 values. At each scale, intraclass correlation coefficients (ICC) (3,1) were then computed between Rest1 and Rest2 for both the raw average BEN values and the derived centile scores.

### Alternative denoising and parcellation

Previous findings suggest that MSE may be influenced by retained signal frequencies (50). To examine whether lifespan trajectories of BEN were affected by temporal filtering and parcellation strategy, MSE was computed from rs-fMRI data using four alternative procedures: 1) the original HCP-preprocessed data without additional temporal filtering; 2) a standard band-pass filter (0.008–0.09 Hz); 3) a stricter band-pass filter (0.03–0.07 Hz); and 4) entropy estimation from ROI-averaged BOLD signals. For this analysis, the Tian S2 subcortical atlas was combined with increasingly finer versions of the Schaefer cortical atlas (100–400 ROIs). After each procedure, mean BEN was used as the dependent variable in the GAMLSS model. For procedure (4), BOLD signals were first averaged within each ROI, entropy was then estimated from the ROI time series (as opposed to computing voxel-wise entropy and then averaging within ROIs), and the resulting ROI-wise entropy values were averaged across regions. For consistency, lifespan trajectories were estimated using the same models described in the primary analyses (see “Modeling normative growth curves across the lifespan” and the SI).

### Correlations with brain metabolism

To test whether MSE is correlated to brain metabolism, cortical maps of CBF, oxygen metabolism, and glucose metabolism (51) were obtained in the fsLR 164k space, down-sampled to 32k, and parcellated using the Schaefer 200 cortical atlas. These maps were derived from a sample of 33 healthy participants aged 20-33 years (51). To match the life stage of this sample, only participants from the HCP-Y dataset were included in this analysis. Regional entropy values at all timescales were previously obtained for each participant (see MSE mapping). Group-level ROI values for each scale were computed by averaging entropy values across participants within each region. Then, Pearson’s correlation was computed between parcel-wise values of each metabolic map and regional entropy values at each timescale. To account for spatial autocorrelation, significance was assessed using spatial autocorrelation-preserving permutation tests (i.e., “spin test”) (5,000 permutations) (52, 53).

### Moderated Mediation

At each entropy scale, a moderated mediation model (39) was tested to determine if the indirect effect of Age (X) on Fluid Intelligence (Y) through average BEN (M) was itself moderated by Age. In this specific formulation, Age serves as both the independent variable and the moderator. For this analysis, we used BEN residuals obtained from the GAMLSS model after removing the effects of gender and site. In addition, age and BEN were mean-centered before the analysis.

The following linear regression models were estimated:

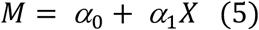

Where *α*_1_ represents the effect of Age on BEN, and

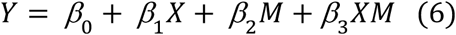

where *β*_2_ represents the effect of BEN on FI and *β*_3_*XM* captures the moderation of the BEN-FI association by age. If the interaction effect was significant, conditional indirect effects (IE)— representing the portion of the age effect on cognition mediated by BEN at any given age—were estimated across the age range of the sample as:

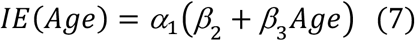

Then, 95% CIs were derived using a nonparametric bootstrap procedure (2,000 resamples).

For each bootstrap resample, both the mediator (Equation 5) and outcome (Equation 6) models were re-estimated. Then, the conditional IE was calculated at any specific age spanning the observed range of the sample (Equation 7).

This procedure produced a bootstrap distribution of the IE at every age. The 95% CIs were determined by the 2.5th and 97.5th percentiles of these distributions, allowing to identify specific developmental windows where the indirect effect was significantly different from zero.

This same approach was applied to all 232 ROIs to map regional-level indirect effects across the lifespan.

### Age prediction

Age prediction was performed using entropy values derived from Rest1. Model performance was evaluated using four algorithms: Ridge Regression (RR), Support Vector Regression (SVR), Random Forest Regression (RFR), Gaussian Process Regression (GPR), and Multilayer Perceptron (MLP).

First, Site- and FD-corrected entropy values from Scale 1 to Scale 3 were averaged for each of the 232 ROIs. Scale 4 and Scale 5 were discarded as preliminary correlation analyses showed high correlation (*r* > 0.9) between Scales 3, 4, and 5, indicating high redundancy between these coarser timescales. Models were trained and tested using age-stratified 10-fold cross-validation (CV). In each iteration, approximately 90% of participants were used for training and 10% were held out for testing. Before each training, feature selection among the 232 ROIs was done using a sparse group lasso (54), whose hyperparameters were identified using grid search on the training set. Hyperparameters of the prediction algorithms were also identified using an inner 5-fold CV. The optimized model was then refit on the full outer training fold and evaluated on the held-out out test set. Model performance was assessed using mean absolute error (MAE) and R^2^ on the held-out test sets.

### Brain Age Gap (BAG)

Using the best performing model, age was predicted for the whole sample. The BAG was then calculated as the difference between predicted and chronological age. Then, the raw BAG was corrected by regressing out the chronological age (55). Finally, Pearson’s correlations were computed between the corrected BAG and fluid intelligence in the whole sample as well as within age bins. To assess the robustness of the results, the correlation was also tested using by calculating the BAG from the age predicted from the out-of-fold test sets BAG during the 10-fold CV.

## Results

### Normative growth curves

At Scale 1, average BEN displayed an increase throughout the lifespan, stabilizing in the late adulthood. After Scale 1, the trajectories exhibit a trend reversal, with BEN decreasing from childhood to late adulthood. At Scale 2, the curve displayed a weak, linear decline with aging. At coarser scales, however, the age-related entropy decline increased and the pattern remained similar across Scales 3, 4, and 5 (Figure 2). Results of bootstrap resampling analysis are included in the SI (Figure S1). Split-half validation demonstrated consistent out-of-sample performance and residual calibration across random partitions of the dataset (Figure S2, Table S1).

**Figure 2.**
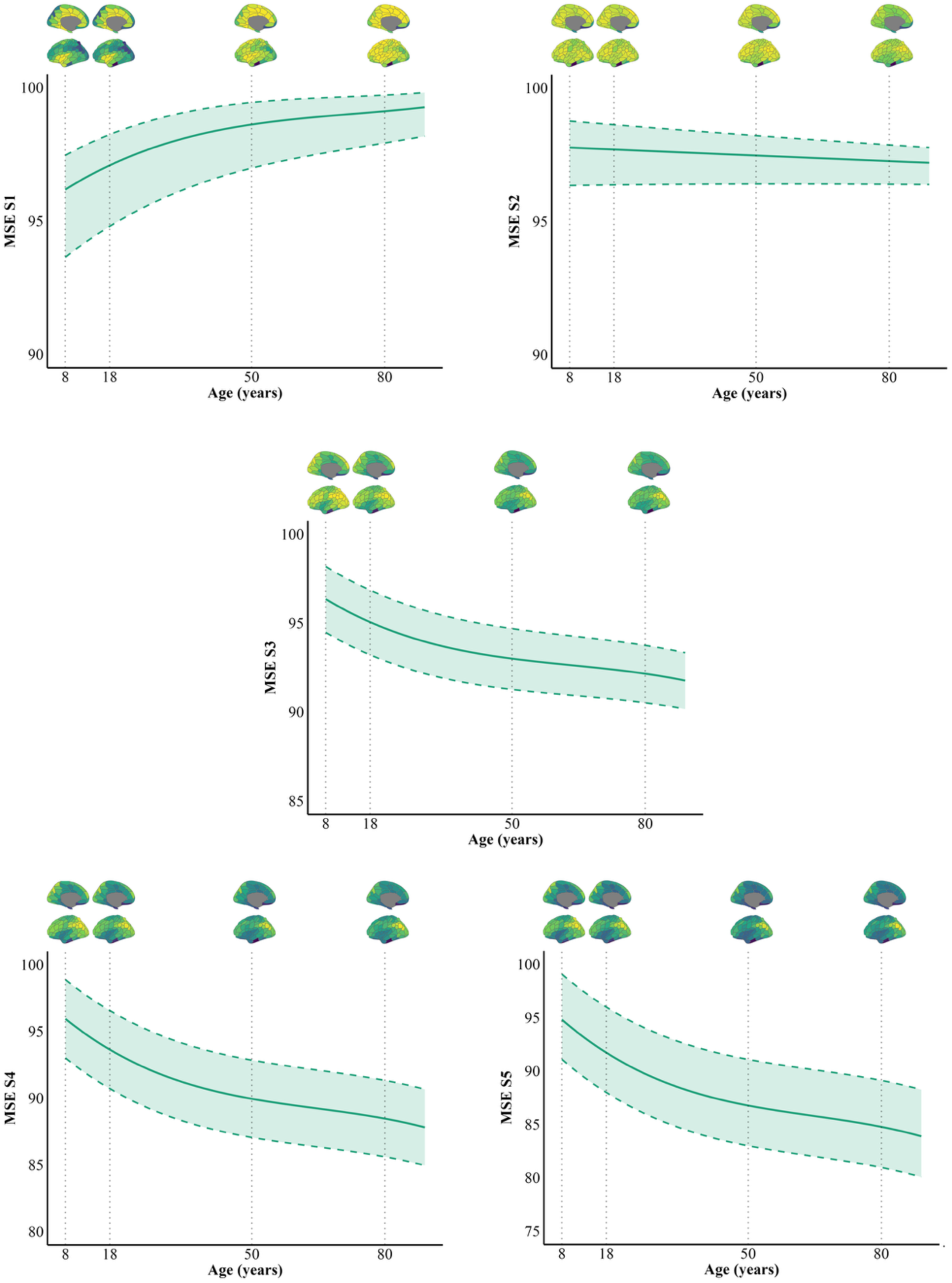
Normative multiscale entropy (MSE) chart across timescales. Average brain entropy (BEN) values across cortical and subcortical areas are normalized to the maximum BEN within each scale. Solid lines represent the median (50th) centile curve, while dashed lines denote the 95% confidence intervals (CI). Note that, due to the different value ranges between timescales, we decided to use a different value limits on the *y* axis. Specifically, the value ranges increase at higher timescales.

### Intraclass correlation

The ICC of the raw BEN between Rest1 and Rest2 ranged between 0.87 and 0.93. The ICC of the centiles ranged between 0.71 and 0.75, with only Scale 2 showing a substantially lower ICC of 0.59 (Figure S3).

### Alternative denoising and parcellation

The same trajectories described above were observed when using MSE estimated from the original HCP preprocessed data, without additional band-pass filtering (Figure S4). Adding a low-pass filter (0.008–0.09 Hz) or applying a stricter band-pass filter (0.03–0.07 Hz), however, disrupted the age trajectories at coarser scales (Figure S5-S6). Calculating entropy from ROI-averaged BOLD signals also influenced the age trajectories, although no differences were observed across parcellations of varying granularity (Figure S7-S10). Specifically, Scale 2 showed an increase across the lifespan, whereas Scale 3 was nearly flat. The trend reversal described in Figure 1 emerged at Scale 4.

### Correlations with brain metabolism

CBF was positively and significantly correlated with BEN at Scale 2 (*r* = 0.33, pspin=0.03) (Figure 3A). Oxygen and Glucose metabolism were positively and significantly correlated with BEN from scale 2 to scale 5, with *r* ranging from 0.33 to 0.46 and from 0.47 to 0.59, respectively. Although not significant, Scale 1 displayed a trend of negative correlation with both Oxygen and Glucose metabolisms (approaching significance for the latter) (Figure 3B-C).

**Figure 3.**
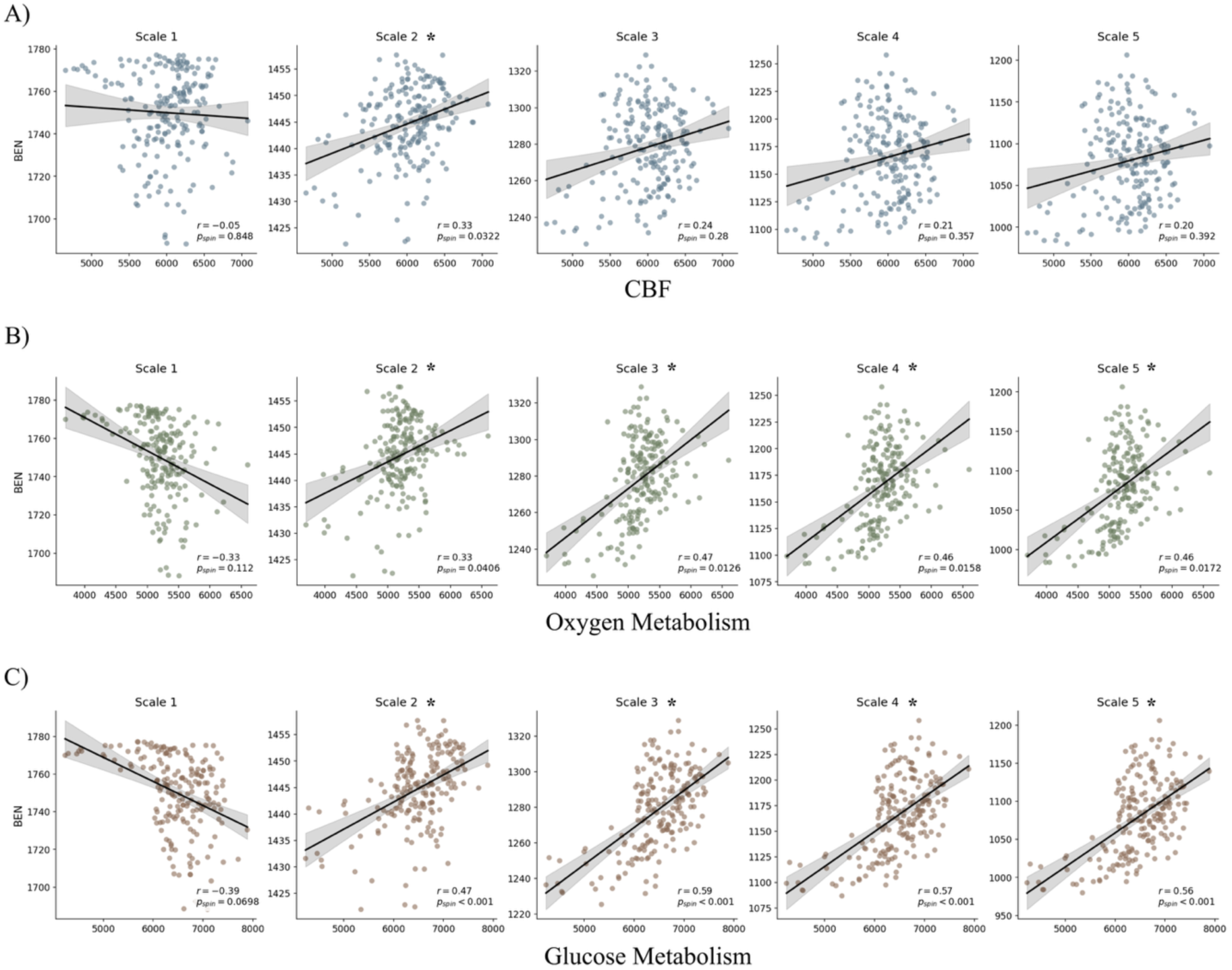
Correlations between brain entropy (BEN) and cerebral blood flow (CBF) (A), oxygen metabolism (B), and glucose metabolism (C). * = significant correlation.

### Moderated mediation

All participants with missing cognitive scores were excluded from the sample (N = 2540). The interaction between average BEN and age was significant for all scales, indicating that the association between BEN and fluid intelligence varies across age. Scale 2 exhibited a substantially attenuated mediation effect, characterized by a smaller effect size, a shifted positive-to-negative transition point, and non-significant positive conditional indirect effects. This likely reflects the near-flat lifespan trajectory of entropy at Scale 2, which represents a transitional timescale between the opposite aging patterns observed at faster and slower scales. Because mediation depends on the association between age and entropy, the attenuated age-related variation at Scale 2 results in a comparatively weak indirect effect. For this reason, Scale 2 results are presented in the SI (Figure S11, Table S2). Across the other significant scales, the indirect effect was positive in early life, became non-significant in midlife, and turned negative in older age (bootstrap 95% CI). The same pattern was observed at the ROIs-level, with stronger IEs in childhood and old age, and non-significant IEs in midlife (Figure 4). At Scale 1, aging leads to higher entropy (α1 = 0.6). At the mean age, higher entropy was associated with higher intelligence (β2 = 0.02). However, the negative interaction (β3 = -0.007) suggests that as entropy gets very high, its positive effect on intelligence starts to reduce. At Scale 3, aging reduces entropy (α1 = -0.7), and at the mean age higher entropy was associated with higher fluid intelligence (β2 = 0.002), such that age-related reductions in entropy contribute to lower fluid intelligence. The positive interaction (β3 = 0.007) implies that for those who manage to keep their entropy high, the benefit to their intelligence is stronger. Scales 4 and 5 showed the steepest decline with age (α1 = -1.17, -1.51). At the mean age, higher entropy was associated with lower fluid intelligence (β2 = -0.006 and -0.01, respectively). The positive interaction (β3 = 0.005 and 0.003) indicates that the negative association between BEN and FI weakens as age increases. To complement this analysis, we performed a secondary moderated mediation analysis (10,000 bootstrap simulations) using a data-driven group partition. For each entropy scale, we defined two age cohorts using a threshold derived from the lifespan trajectory of the IE; specifically, the threshold was set at the chronological point where the sign of the indirect effect transitioned from positive to negative. These age groups were utilized as both the independent variable and the moderator in a formal moderated mediation framework. This approach allowed us to statistically test the robustness of the observed mediation profiles in distinct life stages. Consistent with the previous results, the group-based moderated mediation revealed a significant divergence in the role of brain entropy between cohorts (Table 1).

**Figure 4.**
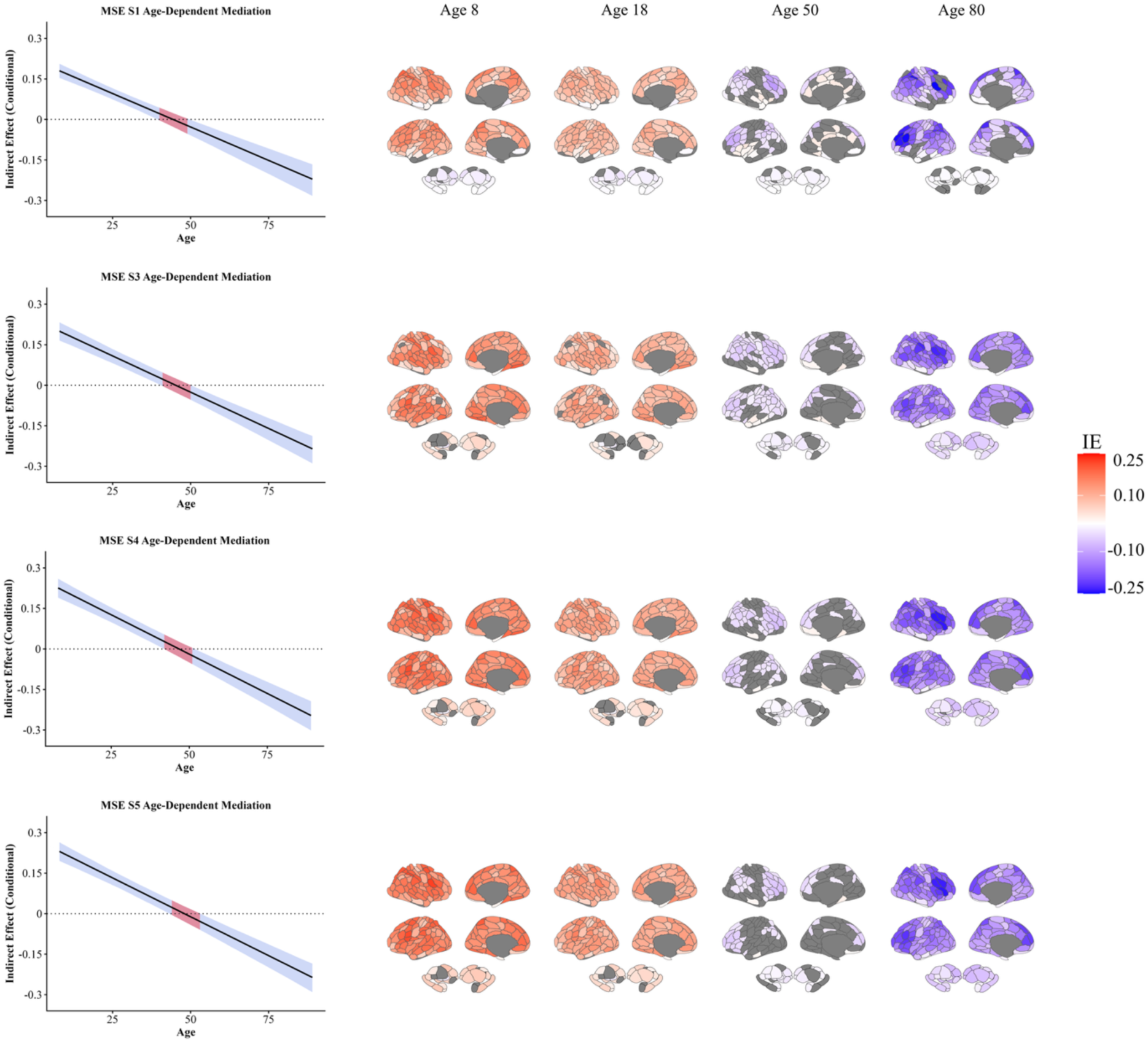
Age-dependent moderated mediation of the relationship between brain entropy (BEN) and fluid intelligence across entropy scales. For each scale, the left panels show the conditional indirect effect of age on fluid intelligence through BEN as a function of age, with shaded areas indicating 95% bootstrap confidence intervals (CI); red segments denote the age range where the CIs include zero. Across scales, the indirect effect is positive in early life, becomes non-significant in midlife, and turns negative in older age. Surface maps illustrate the spatial distribution of region-wise indirect effects at representative ages (8, 18, 50, and 80 years), highlighting a shift from predominantly positive to negative effects across the lifespan. Warmer colors indicate positive indirect effects, whereas cooler colors indicate negative effects. Regions where the CIs of the indirect effect cross zero are depicted in gray.

**Table 1.**
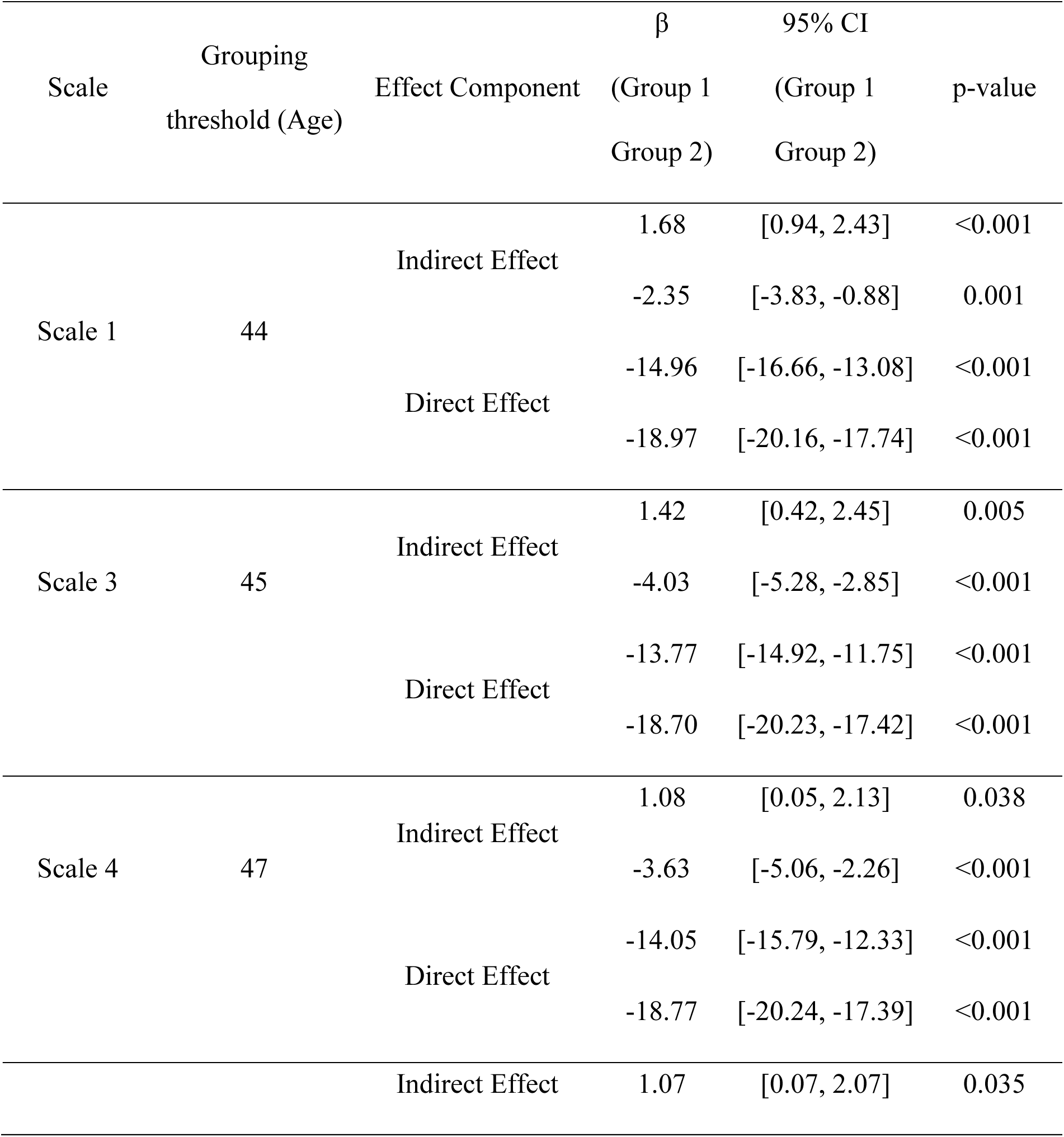
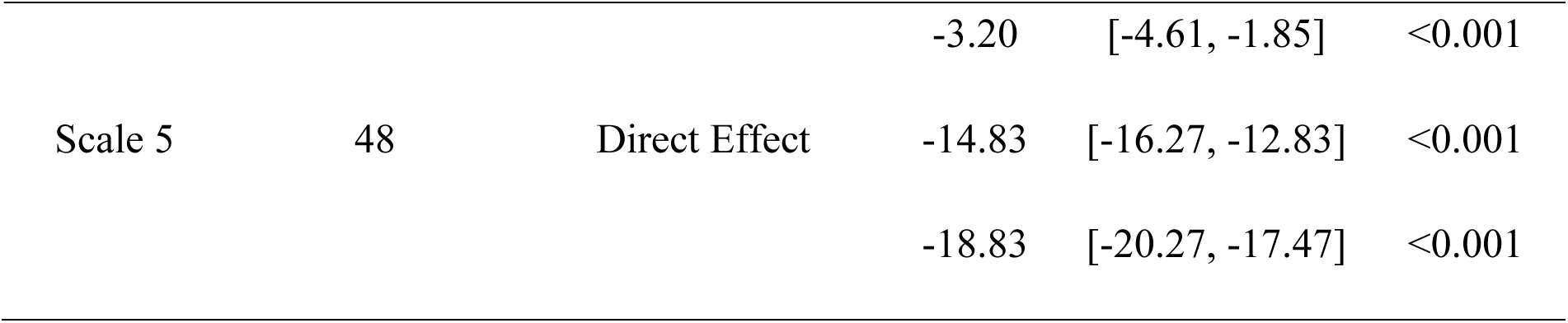
Results of the group-based moderated mediation analysis across entropy scales. For each scale, participants were divided into two age groups using a data-driven threshold corresponding to the age at which the indirect effect (IE) changed sign. β coefficients, 95% confidence intervals (CI), and p-values are reported for both indirect and direct effects in each group. Across scales, the indirect effect differed significantly between the two age-defined groups, supporting an age-dependent change in the mediating role of brain entropy.

### Age prediction

Among the tested models, GPR performed consistently better than other algorithms (R^2^=0.80, MAE=7.11 years), followed by MLP (R^2^=0.78, MAE=7.25 years), SVR (R^2^=0.76, MAE=7.56 years), RR (R^2^=0.74, MAE=8.35 years) and RF (R^2^=0.73, MAE=7.77 years) (Figure 5A). On average, 122.4 ± 5.48 ROIs were retained into the models by the feature selection.

**Figure 5.**
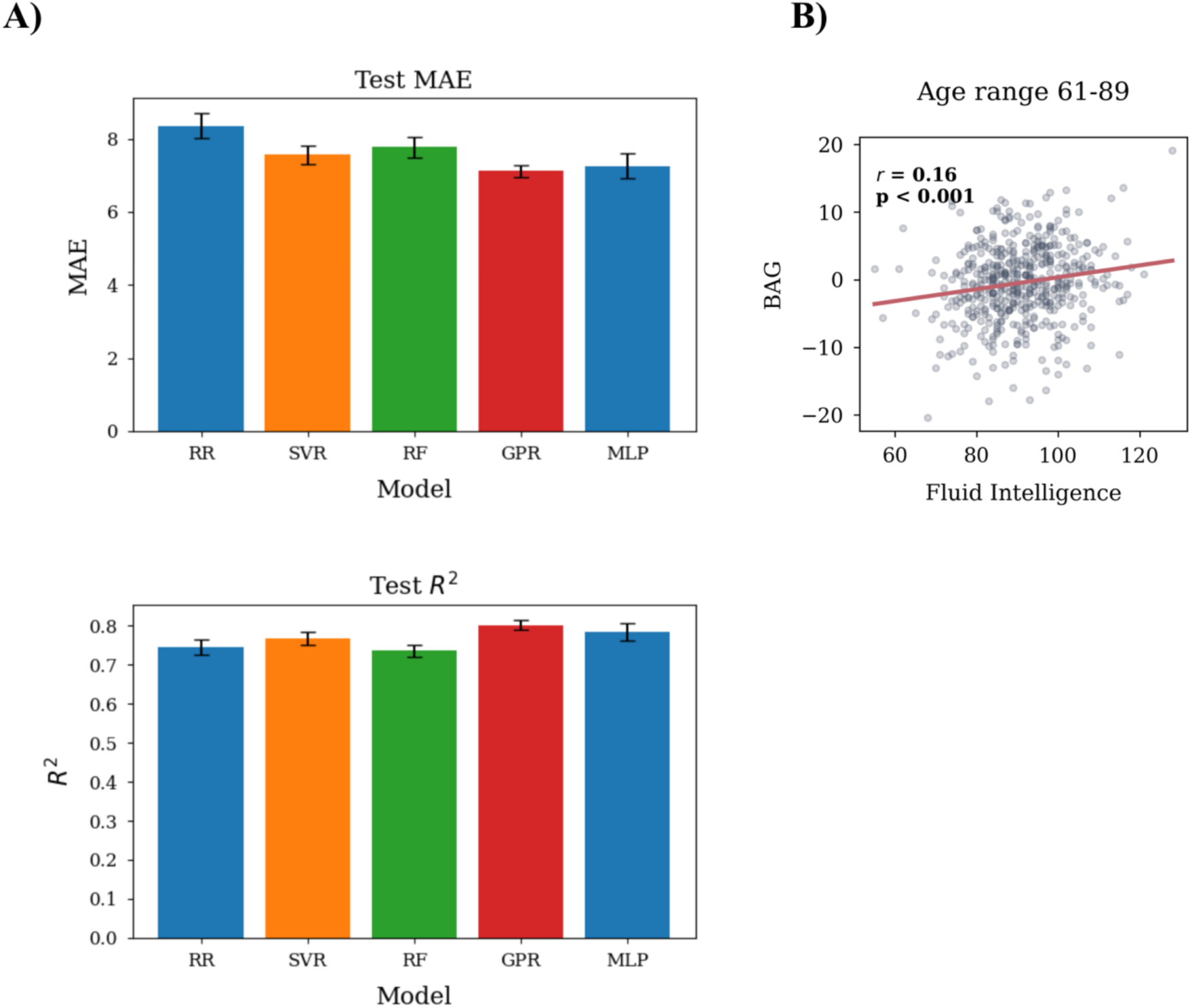
A) Mean absolute error (MAE) and R² are shown across models. Error bars are derived from standard deviation across folds. B) Associations between brain age gap (BAG) and fluid intelligence in participants from 61 to 89 years old. RR = Ridge Regression; SVR = Support Vector Regression; RF = Random Forest; GPR = Gaussian Process Regression; MLP = Multilayer Perceptron.

### BAG

The GPR was used to predict age from the entire sample and calculate the BAG. As demonstrated in previous work (55), the raw BAG was significantly and negatively correlated with chronological age (*r*=−0.38), indicating a systematic overestimation of age in younger participants and underestimation in older participants. This correlation was effectively eliminated after regressing out chronological age (*r*=−0.006; Figure S12). The correlation between BAG and FI in the whole sample was not significant. However, significant correlations emerged when dividing the sample into age bins: 8-30 years, 31-45 years, 46-60 years, and 61-89 years. In the 61-89 years group, BAG was positively correlated to FI (*r*=0.16, p < 0.001), suggesting that an entropy-predicted age higher than the chronological age was related to higher FI scores (Figure 5B). Using BAG calculated (and age-corrected) from the held-out test sets during the 10-CV, this correlation was still significant (*r*=0.18, p < 0.001), and an additional negative and significant correlation emerged in the 31-45 years group (*r*=-0.11, p = 0.004), suggesting that an entropy-predicted age higher than the chronological age was related to lower FI scores (Figure S13).

## Discussion

We used MSE to investigate the developmental and aging trajectories of BEN across different timescales, as well as its associations with brain metabolism and fluid intelligence. These results align with and extended previous findings that showed age-related increase of BEN estimated using SampEn (17, 18, 23). We showed that entropy increases across the lifespan at the finest timescale, before undergoing a trend reversal that begins at scale 2 with a flattening of the trajectory. From scale 3 onward, entropy decreases with age and the pattern does not exhibit meaningful changes at coarser timescales. To our knowledge, this is the first study to examine the lifespan trajectory of BEN across distinct timescales using rs-fMRI data. These findings support the hypothesis that increased entropy at Scale 1 may primarily reflect increased randomness or noise in the signal, whereas signal complexity decreases across the lifespan as captured by coarser timescales (5). In young adults, entropy at Scale 1 was not significantly correlated with metabolic-related measures, albeit displaying a trend of negative correlation that approached significance with glucose metabolism. This pattern reversed at slower scales: CBF was positive correlated with Scale 2, while oxygen and glucose metabolism were positively correlated with entropy from Scale 2 to Scale 5. These results suggest that the complexity of temporal dynamics at these timescales may be linked to underlying metabolic demand and that more complex neural dynamics are energetically costly.

Reliability of entropy was assessed by computing the ICC between raw entropy values calculated from the two rs-fMRI sessions of the HCP. ICC was also computed on centile scores, obtained by mapping Rest1 and Rest2 entropy values to their corresponding positions within a normative distribution estimated using a GAMLSS model fitted on Rest1. Raw entropy showed a good reliability, with ICC ranging from 0.87 at Scale 2 to 0.93 at Scale 1. Centile scores displayed moderate reliability, with ICC ranging from 0.71 at Scale 1 to 0.75 at Scale 4 and 5. Only Scale 2 showed a substantially lower reliability.

We also estimated lifespan trajectories of entropy under alternative temporal filtering strategies. The results showed that the trajectories are strongly influenced by the choice of band-pass filter. In addition, we examined an alternative approach to regional entropy estimation, in which the BOLD signal was first averaged within each ROI and entropy was subsequently computed on the averaged signal (as opposed to computing voxel-wise entropy and then averaging within ROIs). This approach also influenced the observed lifespan trajectories, with the flattening of the curve emerging at scale 3. No differences emerged using finer or coarser parcellations. These findings (29)indicate that entropy estimates derived from MSE are sensitive to the retained signal frequencies and therefore depend on both preprocessing choices and the level at which the signal is summarized. These results align with previous findings (50) and could explain inconsistencies observed between the aging trajectories described herein and in previous works (10). While previous studies have reported a negative association between BEN and FI (18), they did not account for the possibility that entropy reflects distinct neurophysiological processes across lifespan and timescales. Using a moderated mediation framework, we showed that BEN mediates the association between age and FI, and that the relationship between BEN and FI is itself moderated by age. In other words, the direction of the association between entropy and cognition varies across the lifespan, such that higher entropy may be associated with better or worse cognitive performance depending on age. The mediated pathway through BEN indicates that entropy can either attenuate or exacerbate age-related differences in cognitive performance, depending on life stage. Across timescales, this reversal occurs during midlife, when the mediation effect becomes weaker and not significant. Moreover, entropy at faster and slower timescales shows symmetrical effects: at the finest timescale, higher entropy attenuates the negative association between age and cognition earlier in life but exacerbates it in later life, whereas at coarser timescales entropy shows the opposite pattern, exacerbating age-related differences earlier in life and attenuating them in later life.

Finally, we used machine learning to predict age through MSE. The best accuracy was achieved using the GPR (R^2^=0.80, MAE=7.25 years). BAG was weakly associated with FI in an age-dependent manner. In fact, in the 61–89 years age range, BAG was positively correlated with cognition, such that an “older” predicted brain age was associated with higher fluid intelligence. Additional analysis also showed a negative BAG-FI correlation in middle-aged adults (31-45 years), hinting that in this age range that an “older” predicted brain age might be related with lower fluid intelligence. These results further support BEN as a functional biomarker of both aging and cognitive functioning. Specifically, they suggest that lower entropy patterns relative to age-matched individuals may be associated with better cognitive performance during midlife, whereas higher entropy patterns may be associated with better cognition in older age. This study has several limitations that should be addressed in future work. First, it has been shown that the classic MSE algorithm introduces a bias that systematically reduces entropy values as the timescale increases (56). However, the systematic bias was derived assuming a Gaussian process and applies to Gaussian distributed data mostly. As shown in (56), the bias negligibly affects the 1/f process such as rsfMRI time series. Moreover, while this makes the values of entropy at different scales inherently different, our analysis evaluates each timescale independently rather than comparing them. The association between entropy and brain metabolism was examined only within a limited age range. Future studies should extend these findings by investigating these relationships across the lifespan. In addition, entropy-based age prediction was not validated on an independent sample, limiting the generalizability of these results. Similarly, we demonstrated only weak, albeit significant, correlations between BAG and fluid intelligence, and one of these correlations was significant only upon an additional analysis. Therefore, these results need to be further explored and validated in independent samples.

## Conclusions

We used MSE to characterize lifespan trajectories, associations with markers of brain metabolism, and relationships with FI of brain signal temporal dynamics across multiple timescales. We found that faster and slower timescales exhibit distinct and symmetric aging trajectories: entropy increases across the lifespan at Scale 1 and decreases from Scale 3 onward, with Scale 2 marking a transition between these opposing patterns. We propose that increased entropy at faster scales reflects greater randomness or noise in the brain signal, mirrored by a symmetric reduction of complexity at slower timescales. Entropy metrics showed moderate-to-good test–retest reliability. Importantly, lifespan trajectories were highly sensitive to frequency content and to the level of signal summarization, highlighting the need to account for these factors in future studies. Only coarser timescales exhibited positive associations with markers of brain metabolism, suggesting a link between more complex temporal dynamics and energetic demand. Furthermore, BEN mediated the association between age and FI, with the BEN–FI relationship itself moderated by age. Finally, BAG derived from entropy-based age prediction was associated with FI in an age-dependent manner.

## Acknowledgements

This work was supported by the National Institute on Aging [R21AG082345, R01AG070227, R01AG081693, R21AG080518]. The authors declare no competing interests.

