## Supplementary Information for "Lifespan trajectories, metabolic associations, and links to fluid intelligence of brain temporal dynamics characterized by Multiscale Entropy"

### Materials and methods

#### Modeling normative growth curves across the lifespan

##### GAMLSS framework

To estimate normative growth patterns for BEN across cohorts at each scale, Generalized Additive Models for Location, Scale, and Shape (GAMLSS) were applied using the GAMLSS package (version 5.4.22) in R 3.6.3 (22). Model fitting involved selection of an appropriate distribution family and specification of parameter models for the metric of interest.

Among the candidate distributions, the Gumbel (GU) distribution (μ, σ) provided the best fit for Scale 1; Johnson's SU (JSU) (μ, σ, ν, τ) for Scale 2; Skew t Type 4 (ST4) (μ, σ, ν, τ) for Scale 3; t-family (TF) (μ, σ, ν) for Scale 4; and Skew t Type 3 (ST3) (μ, σ, ν, τ) for Scale 5. Nonlinear effects of age were modeled using B-spline functions, denoted as $y = f(Age)$, for Scales 1, 3, 4, and 5, and using a linear term at Scale 2.

Entropy values, denoted by $y$, at each scale were modeled as:

**Scale 1**:

$$y = GU(\mu, \sigma)$$

$$\mu = \beta_0 + f_{df=3}(Age) + \beta_1 Sex + \beta_2 Site + \beta_3 MeanFD \ +$$

$$\sigma = \beta_0 + \beta_1 age + \beta_2 sex + \beta_3 Site$$

**Scale 2**:

$$y = JSU(\mu, \sigma, \nu, \tau)$$

$$\mu = \beta_0 + \beta_1 Age + \beta_2 Sex + \beta_3 Site + \beta_4 MeanFD$$

$$\sigma = \beta_0 + \beta_1 age + \beta_2 sex + \beta_3 Site$$

$\nu = \beta_0$

$\tau = \beta_0$

**Scale 3**:

$y = ST4(\mu, \sigma, \nu, \tau)$

$\mu = \beta_0 + f_{df=3}(Age) + \beta_1 Sex + \beta_2 Site + \ \beta_3 MeanFD$

$\sigma = \beta_0 + \beta_1 age + \beta_2 sex\ \beta + {}_3 Site$

$\nu = \beta_0$

$\tau = \beta_0$

**Scale 4**:

$y = TF(\mu, \sigma, \nu)$

$\mu = \beta_0 + f_{df=3}(Age) + \beta_1 Sex + \beta_2 Site\ \beta + {}_3 MeanFD$

$\sigma = \beta_0 + \beta_1 age + \beta_2 sex\ \beta + {}_3 Site$

$\nu = \beta_0$

**Scale 5**:

$y = ST3(\mu, \sigma, \nu, \tau)$

$\mu = \beta_0 + f_{df=3}(Age) + \beta_1 Sex + \beta_2 Site + \beta_3 MeanFD$

$\sigma = \beta_0 + \beta_1 age + \beta_2 sex\ \beta + {}_3 Site$

$\nu = \beta_0$

$\tau = \beta_0$

### Results

#### Sensitivity analyses

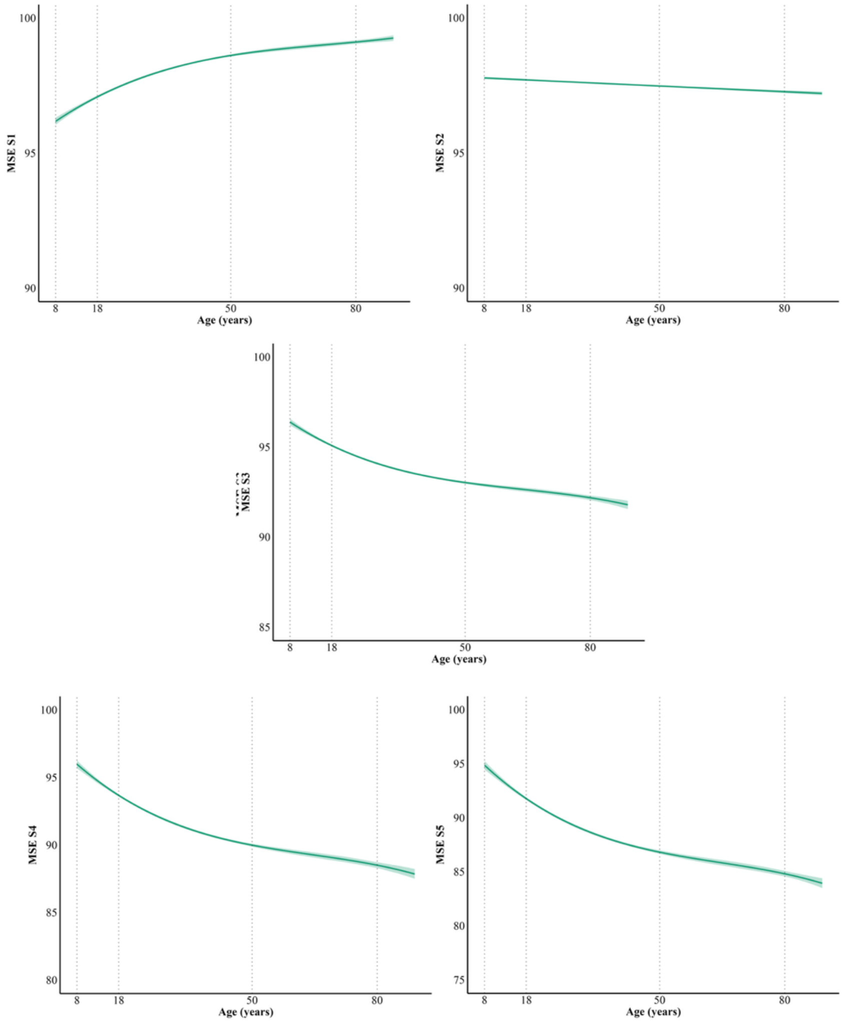


**Figure S1.** 1,000 bootstrap repetitions were conducted with replacement sampling. The sampling preserved the age, sex, and site proportionality of the original cohort by stratifying the lifespan based on ten age intervals (from 8 to 89 years). For each entropy scale, 1,000 bootstrap estimates of the 50th centile growth curve were generated, and 95% bootstrap confidence intervals were calculated from the distribution of bootstrap estimates at each age.

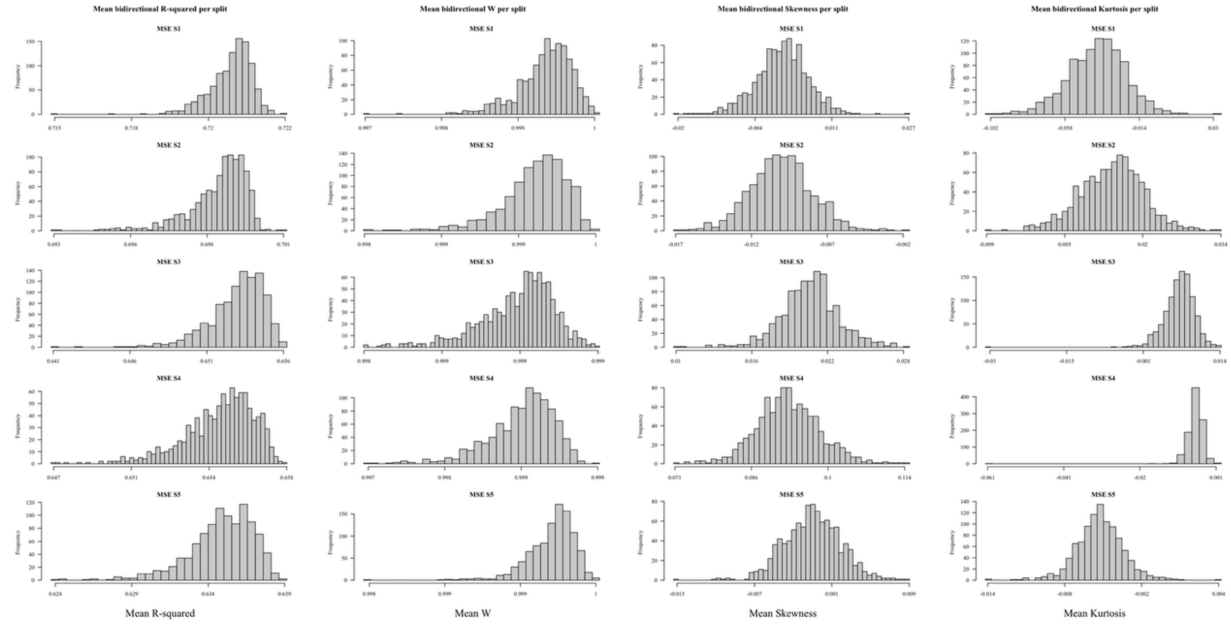


**Figure S2.** Split-half replication analysis across entropy scales. Repeated bidirectional split-half validation (1,000 repetitions) was performed for each entropy scale (Scales 1–5). For each repetition, the dataset was randomly divided into two equal halves stratified by site, and model training and testing were conducted in both directions. Histograms display the distributions of the mean bidirectional out-of-sample R², Shapiro–Wilk statistic (W), residual skewness, and excess kurtosis across the 1,000 repetitions. The narrow distributions of R² and residual diagnostics indicate stable model performance and calibration across random data partitions.

| Scale | R² Median (95% CI) | W Median (95% CI) | Skewness Median (95% CI) | Kurtosis Median (95% CI) |
|---|---|---|---|---|
| Scale 1 | 0.720 (0.719–0.0.721) | 0.9993 (0.9985–0.9994) | 0.001 (-0.009–0.010) | -0.037 (-0.074– -0.006) |
| Scale 2 | 0.699 (0.695–0.700) | 0.9992 (0.9988–0.9994) | -0.009 (-0.013– -0.005) | 0.014 (0.002–0.025) |
| Scale 3 | 0.653 (0.648–0.655) | 0.9990 (0.9985–0.9992) | 0.020 (0.016–0.025) | 0.006 (0.0002–0.010) |

| | | | | |
|---|---|---|---|---|
| Scale 4 | 0.655<br>(0.650–0.656) | 0.9985<br>(0.9980–0.9988) | 0.09<br>(0.08–0.10) | -0.004<br>(-0.008– -0.001) |
| Scale 5 | 0.635<br>(0.629–0.637) | 0.9993<br>(0.9989–0.9995) | -0.0007<br>(-0.006–0.004) | -0.005<br>(-0.009– -0.001) |

**Table S1.** Results of repeated bidirectional split-half validation across entropy scales. Values represent the median and empirical 95% confidence interval (2.5th–97.5th percentiles) across 1,000 repetitions. $R^2$ was calculated on the held-out test set, whereas Shapiro–Wilk W, skewness, and excess kurtosis were derived from quantile randomized residuals of the fitted training models.

#### Intraclass correlation

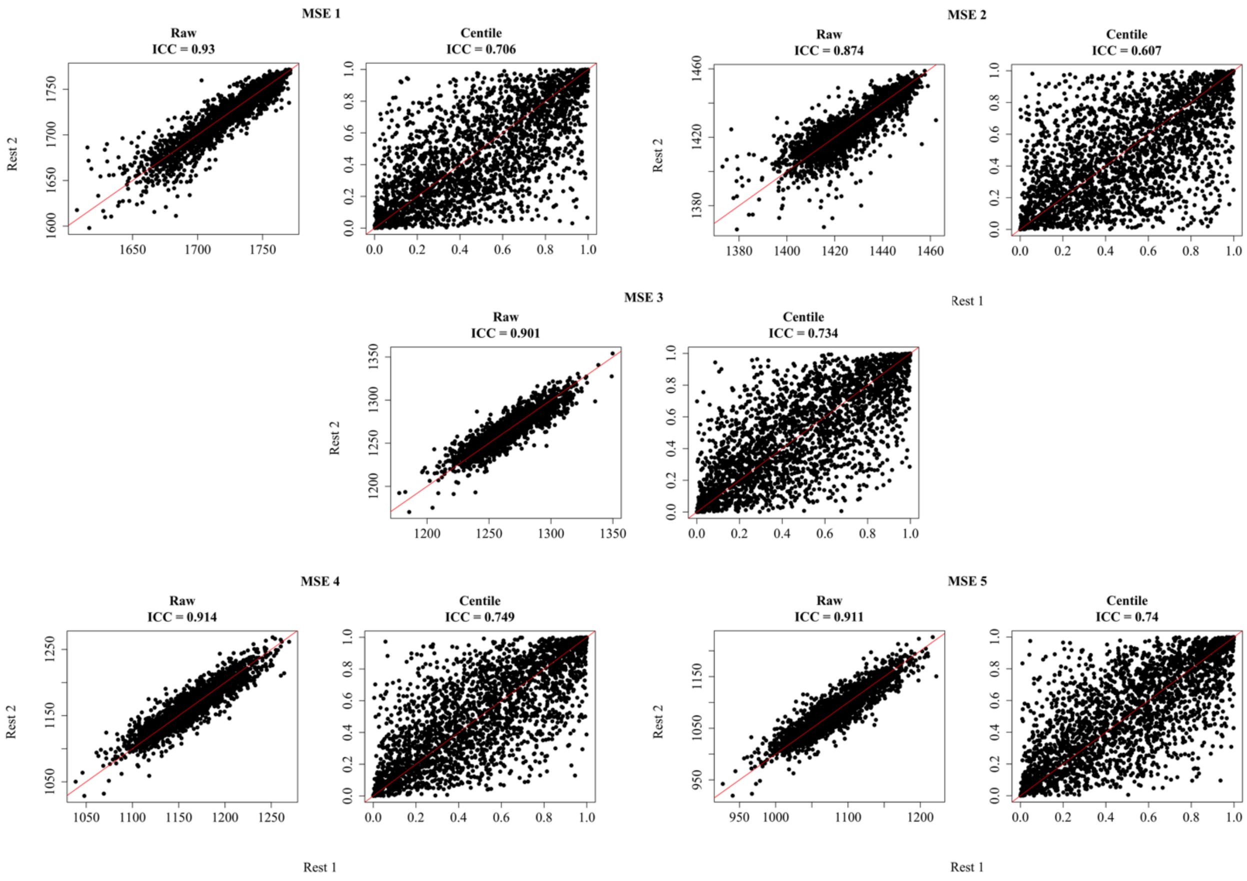


**Figure S3.** Intraclass correlation of Multiscale entropy (MSE) estimated using Rest1 and Rest2 rsfMRI sessions from the Human Connectome Projects (HCP). Plots on the left show the correlation between entropy raw values. Plots on the right show the correlation between participants' centile scores estimated from Rest1 and Rest2. For Rest2, centile scores were estimated using the normative model parameters derived from Rest1.

#### Alternative denoising and parcellation

##### HCP denoising

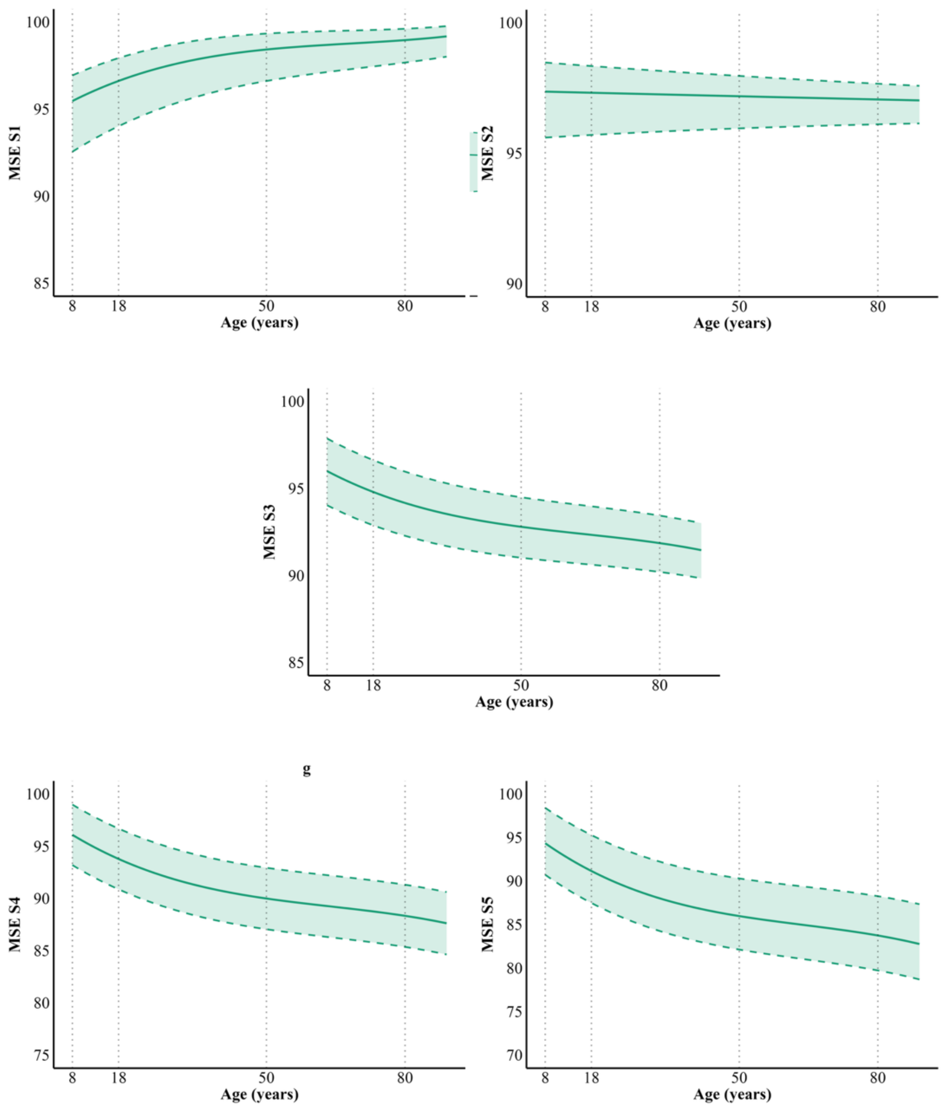


**Figure S4.** Normative multiscale entropy (MSE) chart across timescales. Original preprocessed HCP were used to compute MSE, without applying any additional temporal filtering. Average brain entropy values across cortical and subcortical areas, normalized to the maximum entropy. Average entropy was estimated on the HCP original preprocessed data, without applying any bandpass filter. Solid lines represent mean BEN, while dashed lines denote the 95% confidence intervals (CI).

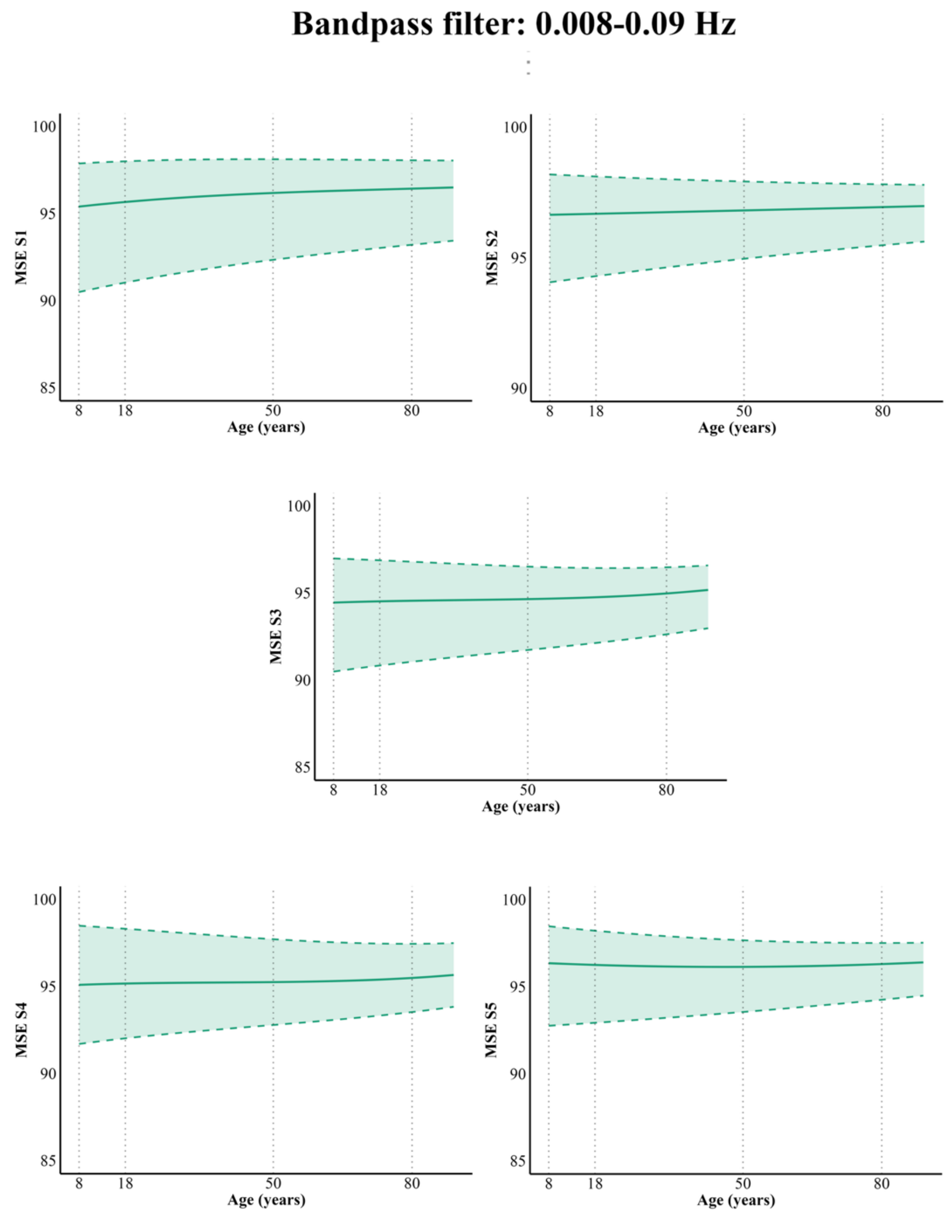


**Figure S5.** Normative multiscale entropy (MSE) chart across timescales. A 0.008-0.09 Hz bandpass filter was applied to the rs-fMRI data before calculating MSE. Average brain entropy values across cortical and subcortical areas, normalized to the maximum entropy. Average entropy was estimated on the HCP original preprocessed data, without applying any bandpass filter. Solid lines represent mean BEN, while dashed lines denote the 95% confidence intervals (CI).

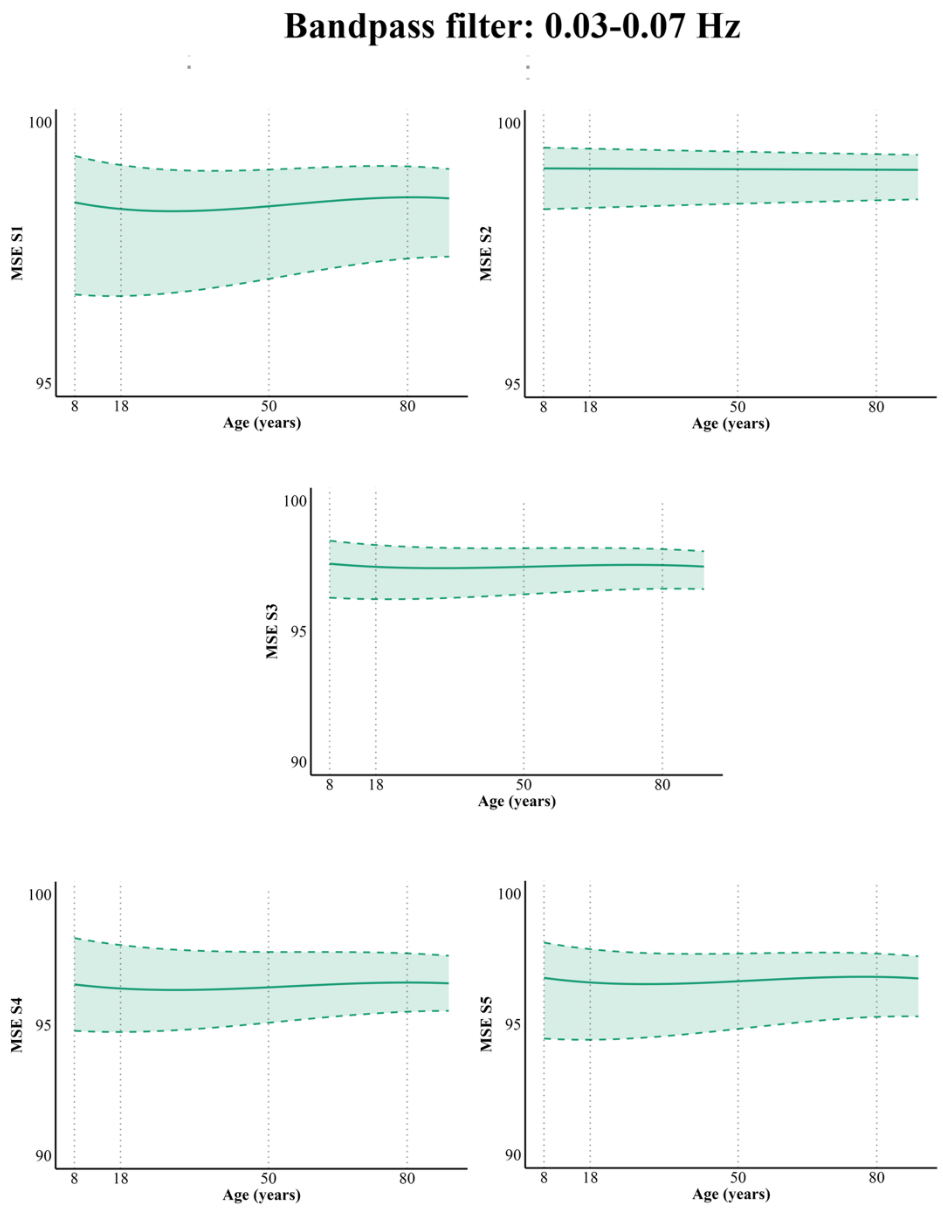


**Figure S6.** Normative multiscale entropy (MSE) chart across timescales. A 0.03-0.07 Hz bandpass filter was applied to the rs-fMRI data before calculating MSE. Average brain entropy values across cortical and subcortical areas, normalized to the maximum entropy. Average entropy was estimated on the HCP original preprocessed data, without applying any bandpass filter. Solid lines represent mean BEN, while dashed lines denote the 95% confidence intervals (CI).

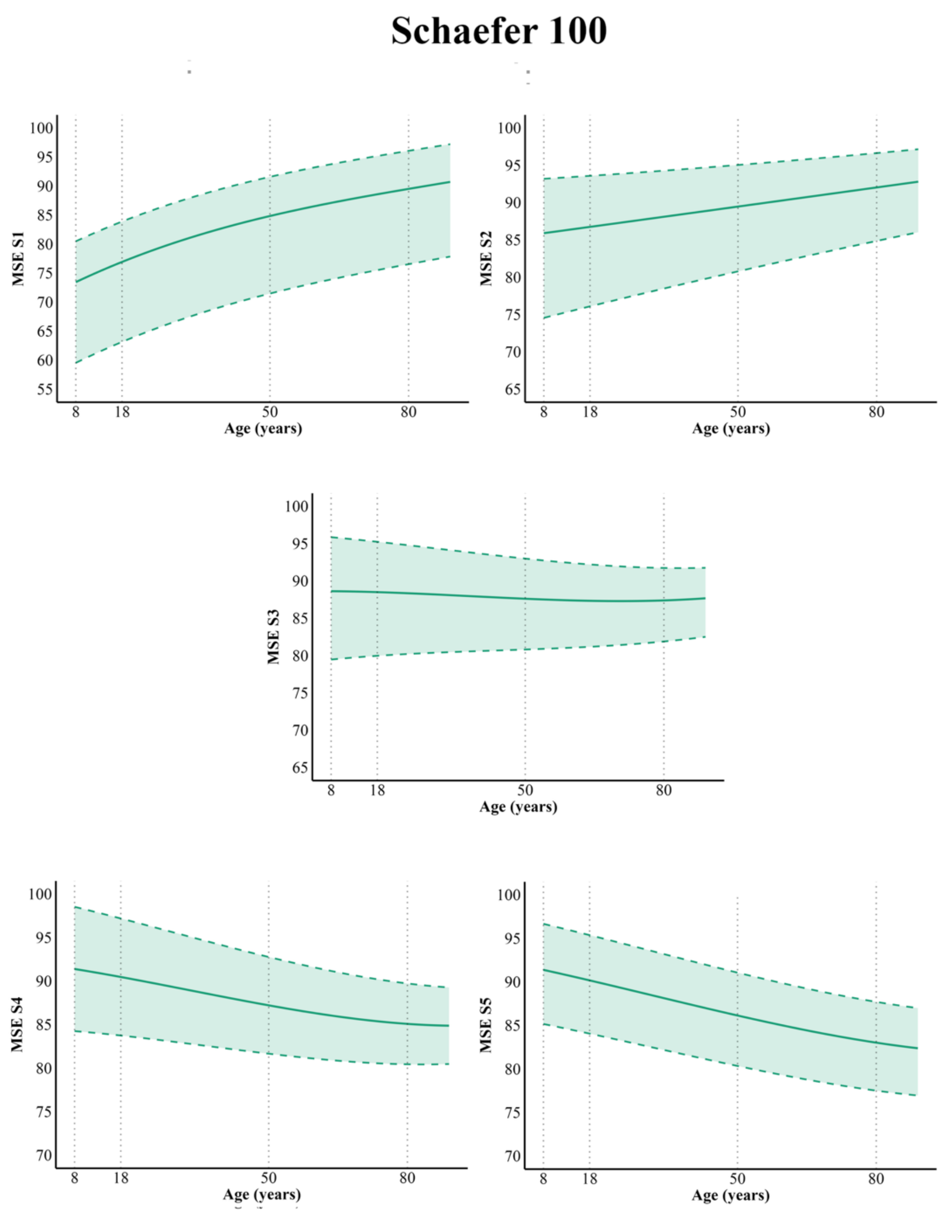


**Figure S7.** Normative multiscale entropy (MSE) chart across timescales. Average brain entropy values across cortical and subcortical areas, normalized to the maximum entropy. Average entropy was estimated by first averaging the BOLD signal within each region of the Schaefer 100 atlas (plus 32 subcortical areas from the Tian S2 atlas) at each timepoint. Entropy was calculated for each region and finally averaged. Solid lines represent mean BEN, while dashed lines denote the 95% confidence intervals (CI).

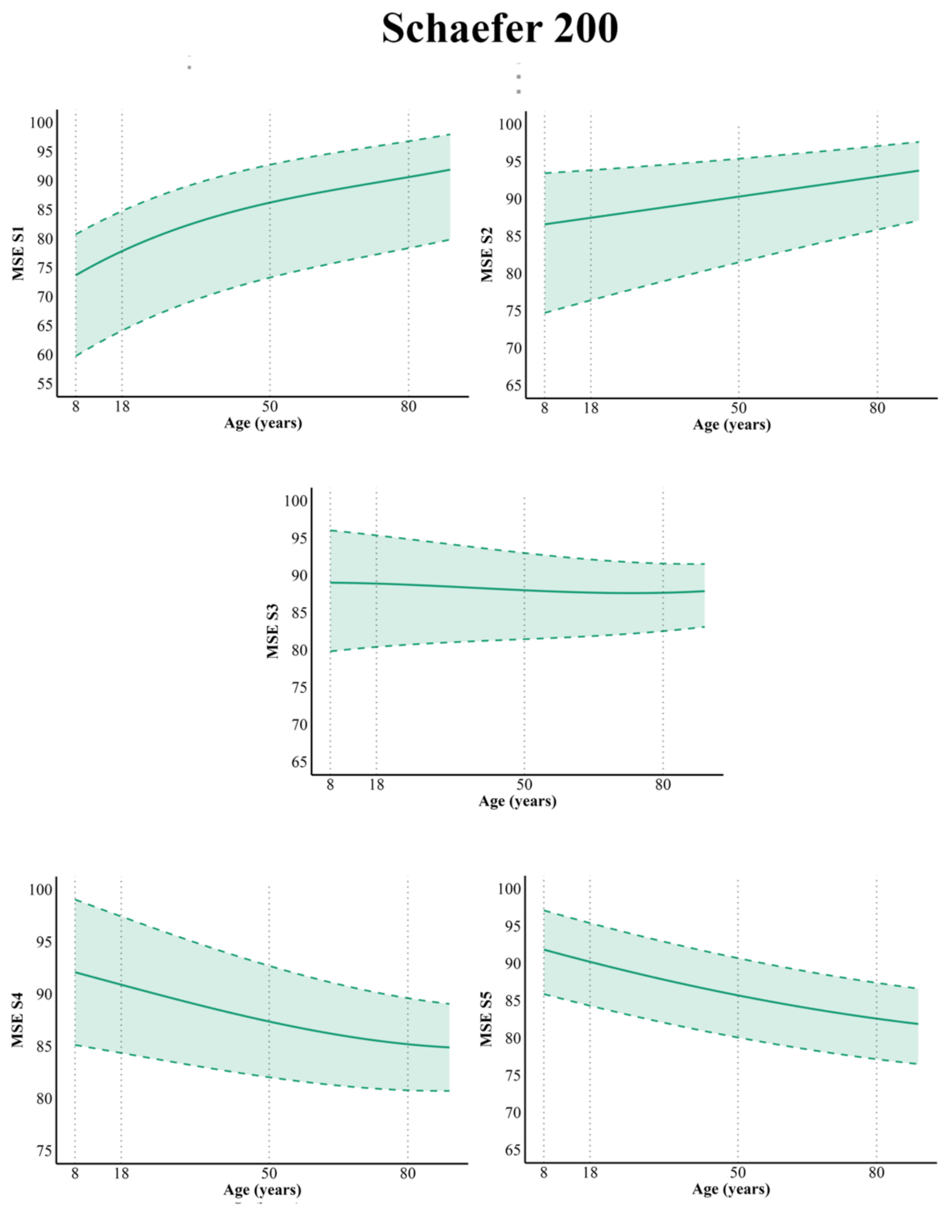


**Figure S8.** Normative multiscale entropy (MSE) chart across timescales. Average brain entropy values across cortical and subcortical areas, normalized to the maximum entropy. Average entropy was estimated by first averaging the BOLD signal within each region of the Schaefer 200 atlas (plus 32 subcortical areas from the Tian S2 atlas) at each timepoint. Entropy was calculated for each region and finally averaged. Solid lines represent mean BEN, while dashed lines denote the 95% confidence intervals (CI).

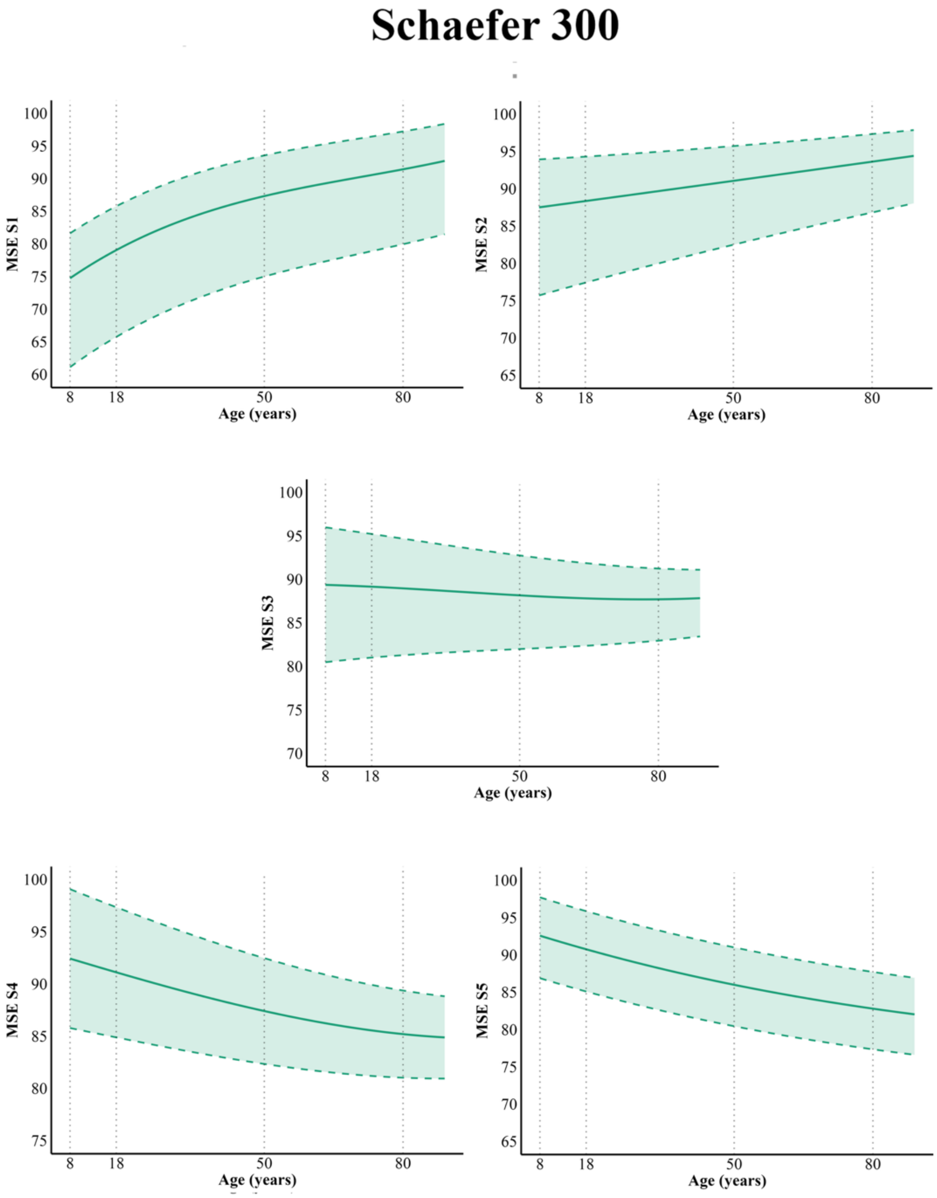


**Figure S9.** Normative multiscale entropy (MSE) chart across timescales. Average brain entropy values across cortical and subcortical areas, normalized to the maximum entropy. Average entropy was estimated by first averaging the BOLD signal within each region of the Schaefer 300 atlas (plus 32 subcortical areas from the Tian S2 atlas) at each timepoint. Entropy was calculated for each region and finally averaged. Solid lines represent mean BEN, while dashed lines denote the 95% confidence intervals (CI).

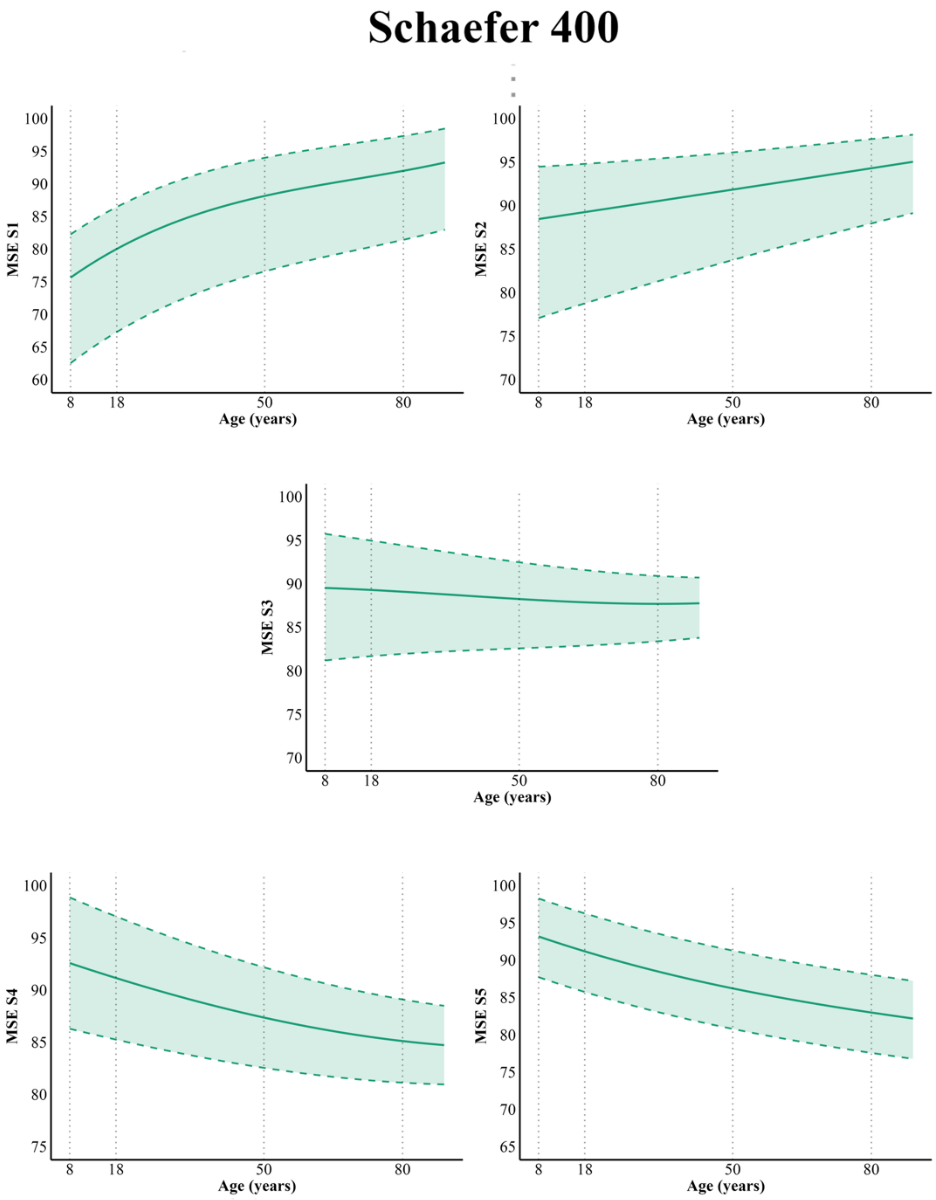


**Figure S10.** Normative multiscale entropy (MSE) chart across timescales. Average brain entropy values across cortical and subcortical areas, normalized to the maximum entropy. Average entropy was estimated by first averaging the BOLD signal within each region of the Schaefer 400 atlas (plus 32 subcortical areas from the Tian S2 atlas) at each timepoint. Entropy was calculated for each region and finally averaged. Solid lines represent mean BEN, while dashed lines denote the 95% confidence intervals (CI).

**Moderated mediation**

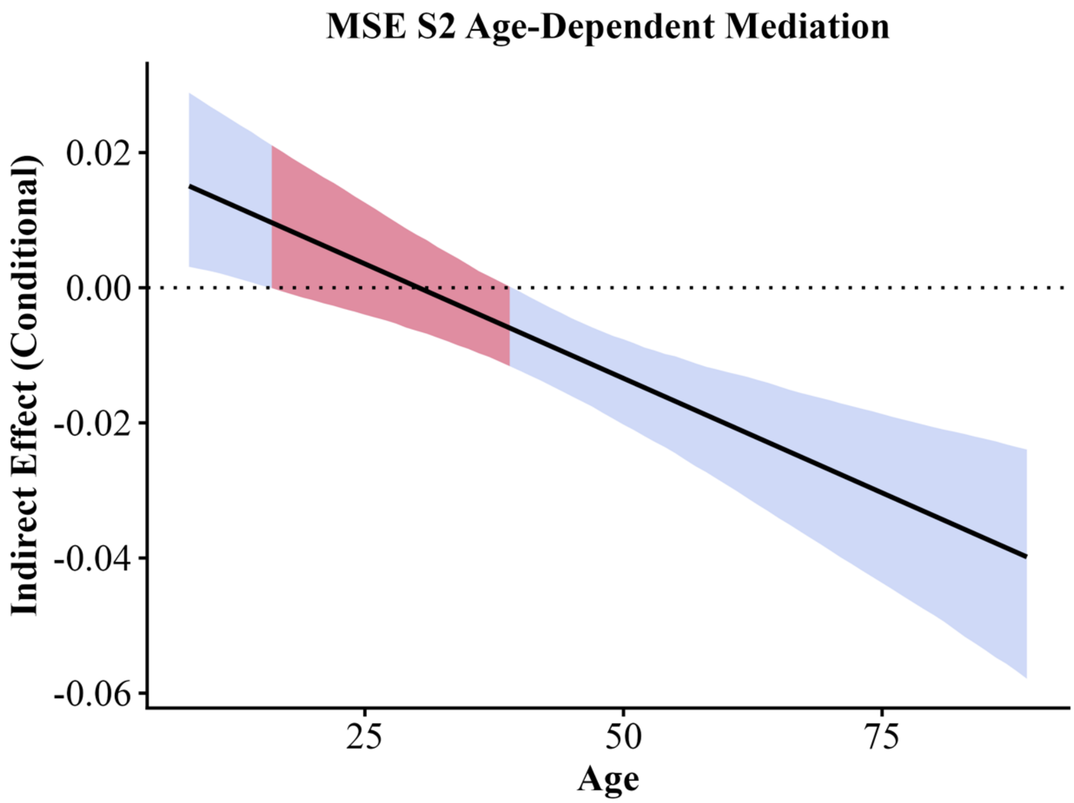


**Figure S11.** Age-dependent moderated mediation of the relationship between brain entropy (BEN) and fluid intelligence at Scale 2. Aging leads to a weak decrease of entropy ($\alpha_1$ = -0.09). At the mean each, entropy has a positive effect on intelligence ($\beta_2$ = 0.06). The positive interaction ($\beta_3$ = 0.006) suggests that the association between entropy and intelligence becomes increasingly positive with age, although entropy itself tends to decrease with aging. The plots shows the conditional indirect effect of age on fluid intelligence through BEN as a function of age, with shaded areas indicating 95% bootstrap confidence intervals (CI); red segments denote the age range where the CIs include zero.

| Scale | Grouping threshold (Age) | Effect Component | β (Group 1 Group 2) | 95% CI (Group 1 Group 2) | p-value |
|---|---|---|---|---|---|
| Scale 2 | 30 | Indirect Effect | 0.28 | [-0.01, 0.6] | 0.03 |
| | | | -1.17 | [-1.58, -0.83] | <0.001 |
| | | Direct Effect | -10.72 | [-11.88, -9.63] | <0.001 |
| | | | -12.18 | [-13.29, -11.15] | <0.001 |

**Table S2.** Results of the group-based moderated mediation analysis at MSE Scale 2. Participants were divided into two age groups using a data-driven threshold corresponding to the age at which the indirect effect (IE) changed sign. β coefficients, 95% confidence intervals (CI), and p-values are reported for both indirect and direct effects in each group. The sign of the indirect effect differed between the two age-defined groups. The indirect effect, however, was significant only in Group 2.

**BAG**

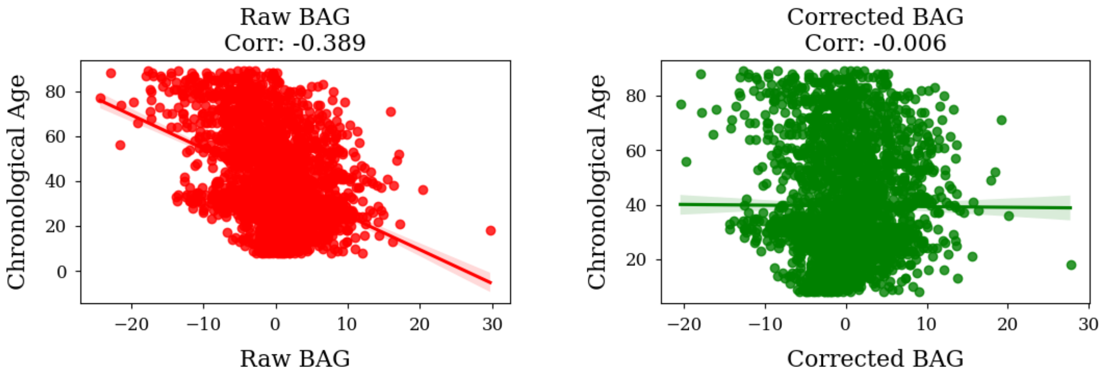


**Figure S12.** Pearson's correlation of chronological age with raw brain age gap (BAG) (left panel) and corrected BAG, after removing the effect of chronological age.

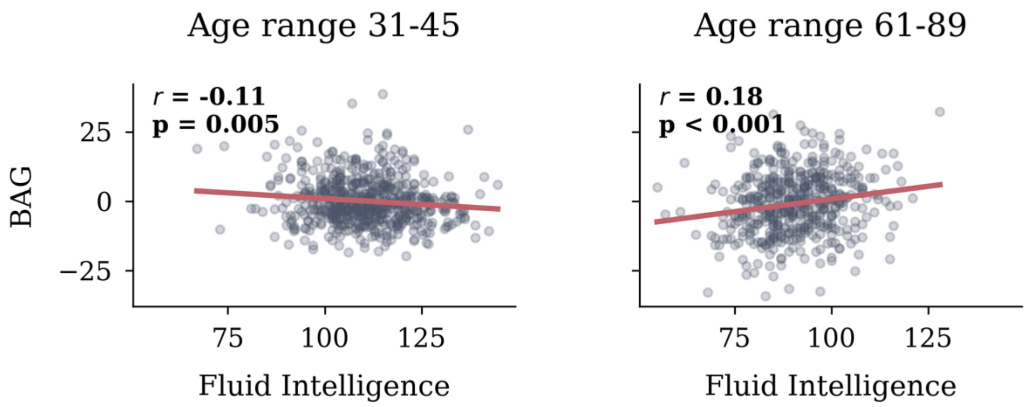


**Figure S13.** Pearson's correlation between brain age gap (BAG) and Fluid intelligence in participants between 31-45 and 61-89 years old. BAG was calculated based on the age predicted in the out-of-fold test sets during 10-fold CV.
